# The full length BEND2 protein is dispensable for spermatogenesis but required for setting the ovarian reserve in mice

**DOI:** 10.1101/2024.02.06.579114

**Authors:** Yan Huang, Nina Bucevic, Carmen Coves, Natalia Felipe-Medina, Marina Marcet-Ortega, Nikoleta Nikou, Cristina Madrid-Sandín, Maria López-Panadés, Carolina Buza, Neus Ferrer Miralles, Antoni Iborra, Anna Pujol, Alberto M. Pendás, Ignasi Roig

## Abstract

Infertility affects up to 12% of couples globally, with genetic factors contributing to nearly half of the cases. Advances in genomic technologies have led to the discovery of genes like *Bend2*, which plays a crucial role in gametogenesis. In the testis, *Bend2* expresses two protein isoforms: full-length and a smaller one. Ablation of both proteins results in an arrested spermatogenesis. Because the *Bend2* locus is on the X chromosome, and the *Bend2^−/y^* mutants are sterile, BEND2’s role in oogenesis remained elusive.

In this study, we employed a novel *Bend2* mutation that blocks the expression of the full-length BEND2 protein but allows the expression of the smaller BEND2 isoform. Interestingly, this mutation does not confer male sterility and mildly affects spermatogenesis. Thus, it allowed us to study the role of BEND2 in oogenesis. Our findings demonstrate that full-length BEND2 is dispensable for male fertility, and its ablation leads to a reduced establishment of the ovarian reserve. These results reveal a critical role for full-length BEND2 in oogenesis and provide insights into the mechanisms underlying the establishment of the ovarian reserve. Furthermore, these findings hold relevance for the diagnostic landscape of human infertility.

## Introduction

Infertility is a complex and multifaceted issue affecting approximately 8-12% of couples worldwide, with genetic factors playing a significant role in up to 50% of cases (Ding and Schimenti, 2021). The intricate process of human reproduction involves a delicate interplay of numerous genes and biological pathways, making the genetic landscape of infertility both diverse and challenging to unravel.

Advancements in genomic technologies and high-throughput sequencing have facilitated the identification of novel genes implicated in reproductive processes. Among these discoveries, *Bend2* has recently emerged as a novel gene involved in mammalian gametogenesis (Malcher et al., 2022). BEND2 is a member of the BEN domain-containing protein family, whose members play diverse roles in cellular processes such as transcriptional regulation, chromatin remodeling, and protein-protein interactions (Abhiman et al., 2008).

Mouse models have been instrumental in identifying novel genes involved in mammalian fertility. This approach has significantly expanded our understanding of the genetic factors underlying reproductive processes in both males and females (Garretson et al., 2023). A recent study reported that BEND2 is required to complete spermatogenesis since it is a crucial regulator of mammalian meiosis in spermatocytes (Ma et al., 2022)Click or tap here to enter text.. BEND2 is specifically expressed in spermatogenic cells shortly before and during the prophase of meiosis I. In the testis, *Bend2* expresses two protein isoforms: full-length, named p140, and smaller one, p80. BEND2 is essential for the transition from zygonema to pachynema, as its knockout in male mice arrests meiosis at this stage. Its absence leads to disrupted synapsis and induced non-homologous chromosomal pairing. BEND2 interacts with multiple chromatin-associated proteins, including components of transcription-repressor complexes. Additionally, BEND2 regulates chromatin accessibility and modifies H3K4me3, influencing the overall chromatin state during spermatogenesis. These functions collectively establish BEND2 as a key regulator of meiosis, gene expression, and chromatin state in mouse male germ cells. Nonetheless, its role in oogenesis remains unexplored since the *Bend2* locus is on the X chromosome, and mutant males are sterile.

In this study, we aimed to elucidate the role of *Bend2* in oogenesis using a novel mutation of *Bend2* that does not confer male sterility. Using this novel *Bend2* mutation, we have investigated the effects of full-length BEND2 ablation on meiosis, oocyte quality, and follicular dynamics. Furthermore, as this mutation does not cause male infertility, it allowed us to investigate the role of BEND2 in mammalian meiosis in more detail. Our findings contribute to expanding our understanding of the genetic factors influencing mammalian fertility and provide potential avenues for diagnosing genetic determinants of human infertility.

Click or tap here to enter text.Click or tap here to enter text.Click or tap here to enter text.Click or tap here to enter text.

## Results

### BEND2 is highly expressed in spermatogenic nuclei before meiosis initiation and during early meiotic prophase I, independent of meiotic recombination

Taking advantage of several unannotated transcripts detected in 14 dpp wild-type mouse testis from our previous RNA sequencing analysis (Marcet Ortega, Roig and Universitat Autònoma de Barcelona. Departament de Biologia Cel·lular, 2016), we found a novel splice variant of the then annotated *Gm15262* gene, which is an X-linked gene containing two BEN domains (**Fig. 1A**). This variant is specifically expressed in mouse testes and fetal ovaries containing germ cells undergoing meiotic prophase I and shows a nuclear localization at the heterochromatin of spermatocytes when the GFP-tagged splice variant is electroporated to the testes of young mice (**Fig S1 A-B**). Thus, we hypothesize this gene could be essential for spermatocyte and oocyte development. Based on sequence homology, we named this *Gm15262* gene as the mouse homolog of the human *Bend2*, and thus, we renamed it *Bend2*, as also proposed by others recently (Ma et al., 2022).

**Figure 1.**
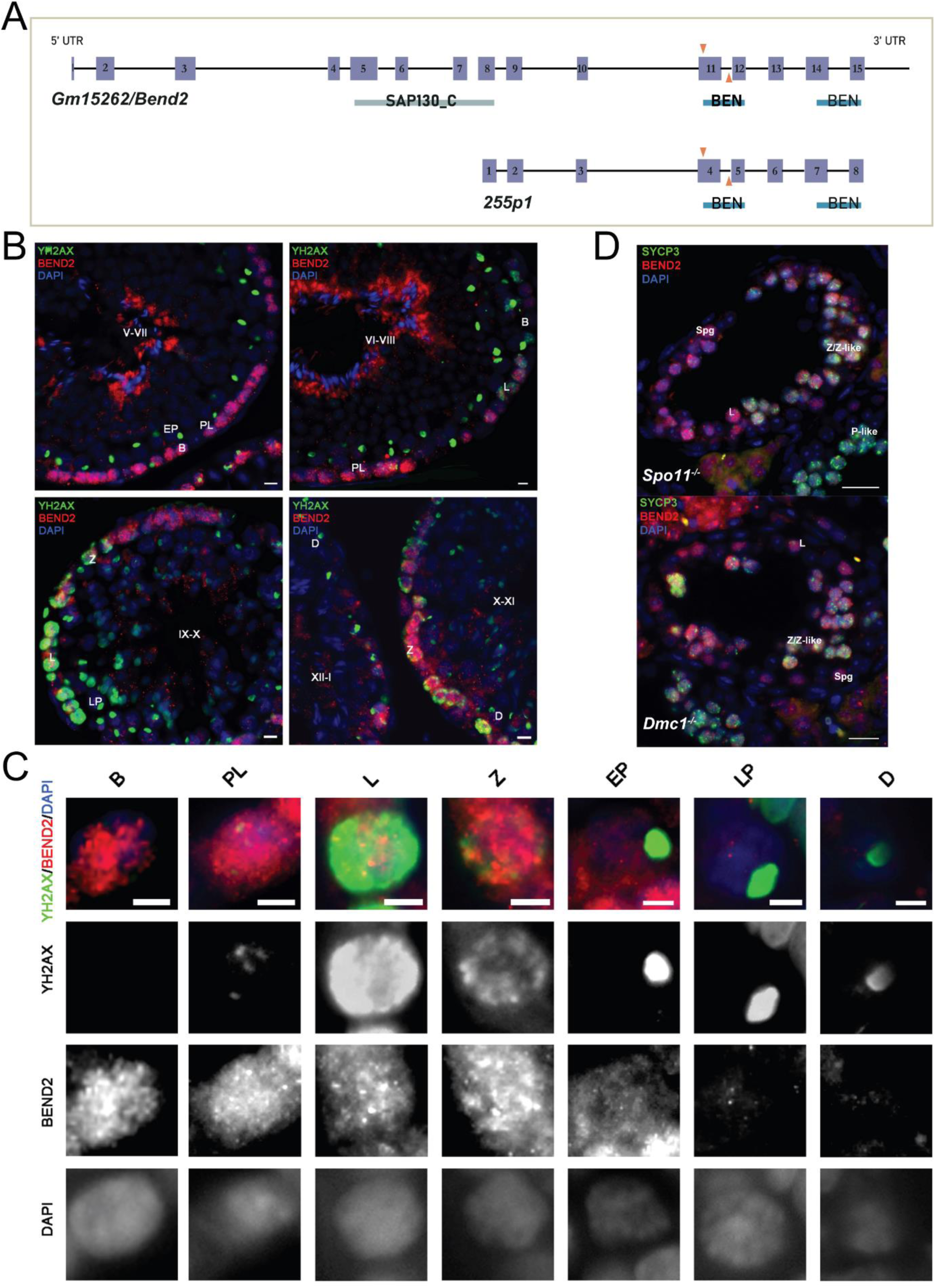
Expression of BEND2 during spermatogenesis. (A) Schematic representation of mouse *Gm15262*/*Bend2* and its novel splice variant *255p1*. The exons are shown as purple boxes. The predicted domains are labeled below the exons. SAP130_C: C-terminal domain of histone deacetylase complex subunit SAP130; BEN: BEN domain. A pair of gRNAs (orange arrows) target exon 11 of *Bend2* within the first BEN domain to generate the *Bend2^Δ11^* mutation by CRISPR/Cas9. The *in-house* BEND2 antibody is generated using full-length 255P1 sequence as immunogen. (B) BEND2 localization in wild-type mouse testis. Staging of the seminiferous epithelium is based on the localization of spermatocytes (indicated by the expression and localization of ϒH2AX(green)) and spermatids (indicated by DAPI). The tubule stage is shown in uppercase Roman numerals. Scale bar, 20 µm. (C) Magnification of BEND2-positive cells from testis sections. Expression of BEND2 was characterized in cells along spermatogenesis from spermatogonia to late diplotene spermatocyte. Scale bar, 10 µm. (D) BEND2 localization in SPO11- and DMC1-deficient testis. In these cases, SYCP3 (green) was used to identify spermatocytes. Scale bar, 50 µm. Spg: spermatogonia; B: B-type spermatogonia; PL: pre-leptotene spermatocyte; L: leptotene spermatocyte; Z: zygotene spermatocyte; Z-like: zygotene-like spermatocyte; EP: early pachytene spermatocyte; LP: late pachytene spermatocyte; P-like: pachytene-like spermatocyte; D: diplotene spermatocyte.

We generated a polyclonal antibody against BEND2 using the novel splice variant of *Bend2* as an immunogen (see validation in **Fig. 2**). To examine BEND2 expression during spermatogenesis, we immunostained PFA-fixed testis sections against BEND2 and the DNA damage marker γH2AX to allow the identification of spermatocytes. BEND2 was abundantly detected in the nuclei of the peripheral cells of tubules from stage V until XII-I (**Fig. 1B).** This nuclear staining first appeared in spermatogonia before DSBs were generated. BEND2 staining was highly present in spermatocytes from pre-leptotene when meiosis initiates and persisted until early pachytene. In late pachytene and diplotene cells, little BEND2 remained in the nuclei of spermatocytes (**Fig. 1C)**. Notably, this expression pattern of BEND2 is reminiscent of the one reported before (Ma et al., 2022).

**Figure 2.**
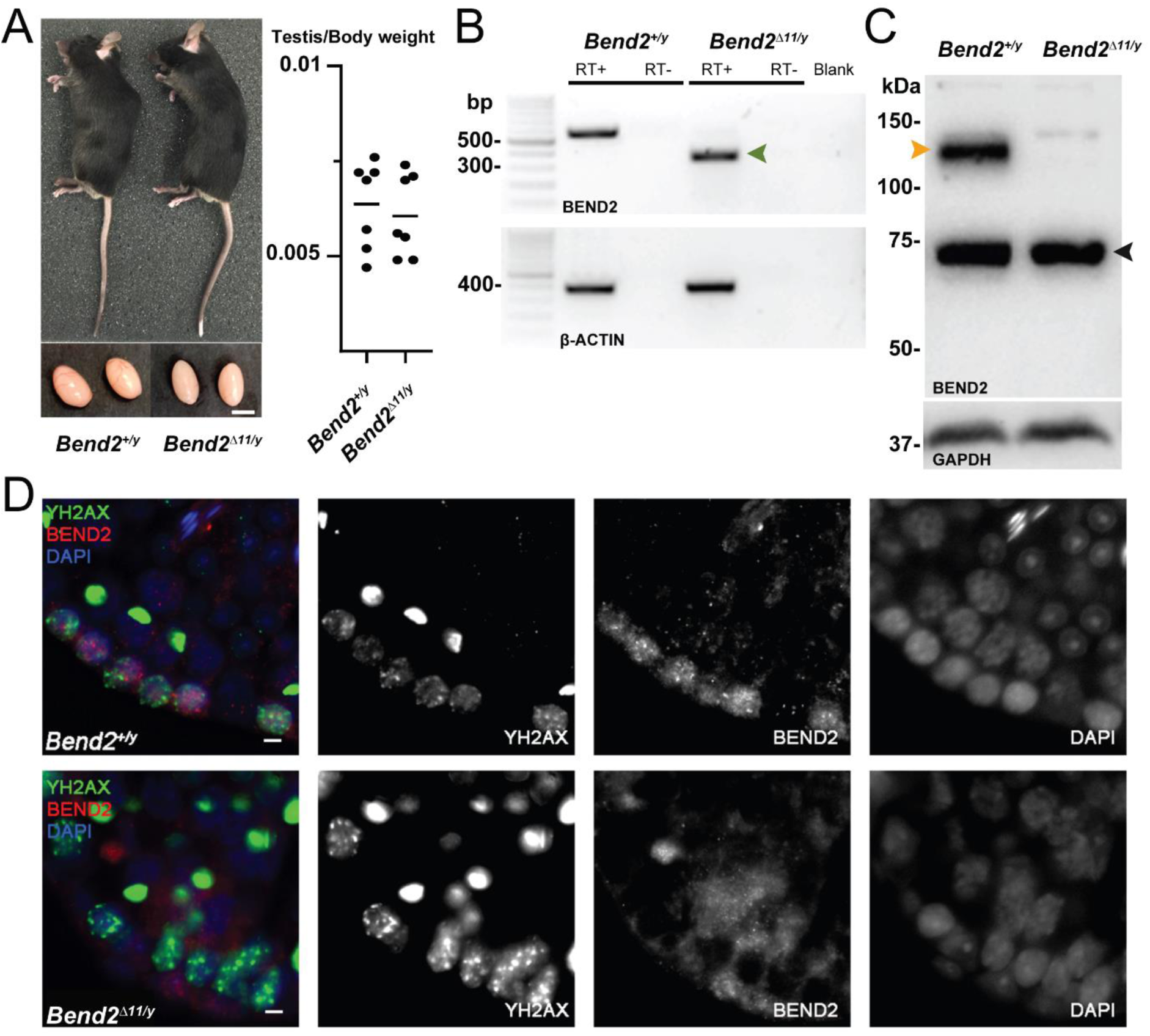
Disruption of BEND2 in *Bend2^Δ11/y^* mice. (A) Mouse appearance and testis size. Scale bar, 5mm. (B) *Bend2* expression in testis by RT-PCR. The expected size of the WT allele was 576bp. Sanger sequencing result showed that the amplified DNA in the mutated allele (372bp, green arrowhead) was from an mRNA that resulted from skipping exon 11 of *Bend2*. (C) Detection of BEND2 by WB from testis protein extracts. The orange arrowhead indicates the full-length BEND2 protein band. An extra protein band ∼75 kDa (black arrowhead) was also detected in both wild-type and mutant testes. (D) Detection of BEND2 by IF. Testis sections were treated with antigen retrieval using Tris-EDTA buffer before staining against BEND2 (red) and SYCP3 (green). The BEND2 staining observed in the *Bend2* mutants closely resembles background staining (not shown), suggesting that the cytoplasmic signal might be nonspecific. Scale bar, 10 µm.

Furthermore, to reveal if the localization of BEND2 in germ cells could have originated as a response to critical events of the meiotic prophase, we examined BEND2 expression in testis sections from recombination-defective mice lacking SPO11. In SPO11-deficient testis, where no DSBs are formed at the onset of meiosis, spermatocytes cannot initiate meiotic recombination, thereby failing to synapse and entering apoptosis (Barchi et al., 2005; Baudat et al., 2000; Pacheco et al., 2015). Interestingly, we found BEND2 was extensively present in nuclei of spermatogonia, leptotene, zygotene, and zygotene-like spermatocytes, as in wild-type mice (**Fig. 1D).** Very occasionally, few BEND2 signals could be observed in some SPO11-deficient spermatocytes, presumably at a more advanced pachytene-like stage. This is consistent with its reduced expression after early pachytene in wild-type mice. Similarly, BEND2 expression was not altered in *Dmc1* mutant spermatocytes (**Fig. 1D)**, which cannot complete meiotic recombination (Barchi et al., 2005; Pittman et al., 1998). Altogether, these results indicate that although BEND2 is highly expressed during early meiotic prophase I when meiotic recombination initiates and progresses, its expression starts as early as in spermatogonia before meiosis begins and is independent of either DSB formation or completion of recombination.

We also examined BEND2 expression in 16 dpc fetal ovary sections by IF and found it localizing at nuclei of some zygotene stage oocytes (**Fig S1C**). However, due to scarce samples and technical obstacles, we could not characterize BEND2 expression in oocytes at earlier stages from 12-14 dpc ovary sections.

### Full-length BEND2 deficiency causes increased apoptosis and persistent unrepaired DSBs in spermatocytes

To address the germ cell functions of BEND2 in mice, we generated BEND2-deficient mice by CRISPR/Cas9. Part of exon 11 of *Bend2* was targeted to be removed to disrupt the first BEN domain (**Fig 1A**). *Bend2^Δ11/y^* mice developed into adults without apparent differences in general physical appearance compared to their littermates. Male *Bend2^Δ11/y^* mice were fertile. The size and weight of *Bend2^Δ11/y^* testes were comparable to that of wild-type testes **(Fig 2A and Table S1**). Similarly to Ma et al., (2022), using our BEND2 antibody, we detected two BEND2-related proteins in wild-type testes by WB: one was ∼130 kDa, and another was ∼75 kDa (**Fig. 2C**). Although the predicted size of full-length BEND2 is 80.9 kDa, based on previous reports (Ma et al., 2022), the 130 kDa protein corresponds to the full-length BEND2, which displays a slower electrophoretic mobility due to its unusual sequence/structure. The 75 kDa protein corresponds to a smaller version of BEND2 lacking the N-terminus, p80 (Ma et al., 2022). Interestingly, we did not detect any BEND2 protein in wild-type testes of 48.3 kDa, which would be expected from the novel transcript 255p1 we identified.

We were unable to detect the full-length BEND2 in *Bend2^Δ11/y^* mouse testis extracts by WB (**Fig. 2C)**. The staining of BEND2 observed in wild-type cells was not present in mutant cells (**Fig. 2D**). However, the p80 BEND2-related protein was still present in *Bend2^Δ11/y^* mutant testes (**Fig. 2C)**. Moreover, an alternative *Bend2* transcript skipping exon 11 was detected in *Bend2^Δ11/y^* testis by RT-PCR and sequencing (**Fig. 2B)**.

We attempted to test if this exon 11-skipped transcript was also present in wild-type mouse testis by RT-PCR and if it represented the p80 BEND2-related protein detected in both wild-type and *Bend2^Δ11/y^* testis by immunoprecipitation (IP) coupled to peptide mass fingerprinting analysis. Unfortunately, only the full-length *Bend2*-202 transcript was consistently amplified in our RT-PCR analysis, and our *in-house* BEND2 antibody did not give satisfactory IP results (*Data was not shown*). Overall, these results suggest that the full-length BEND2 was successfully eliminated from mutant mouse testis. Also, the p80 BEND2 protein, likely corresponding to an exon 11, skipped transcript-is present and might be functional in our mutant testis, based on the observed phenotype (see below).

Histological analysis of *Bend2^Δ11/y^* testes revealed the presence of cells at all the stages of spermatogenesis (**Fig 3A**). However, a significant increase of apoptotic cells was detected in *Bend2^Δ11/y^* testes via TUNEL assays (*p*<0.0001 t-test, **Fig 3B-C**). Further staging analysis showed that *Bend2^Δ11/y^* mice presented a comparable number of apoptotic spermatocytes (66.8 ± 9.7%, mean ± SD, N=4) than wild-type mice (52.5 ± 9.5%, mean ± SD, N=4, *p*>0.05 One-Way ANOVA, **Fig 3D**). Therefore, the full-length BEND2 might be dispensable for completing spermatogenesis, and its absence only caused a slight defect in spermatogenesis.

**Figure 3.**
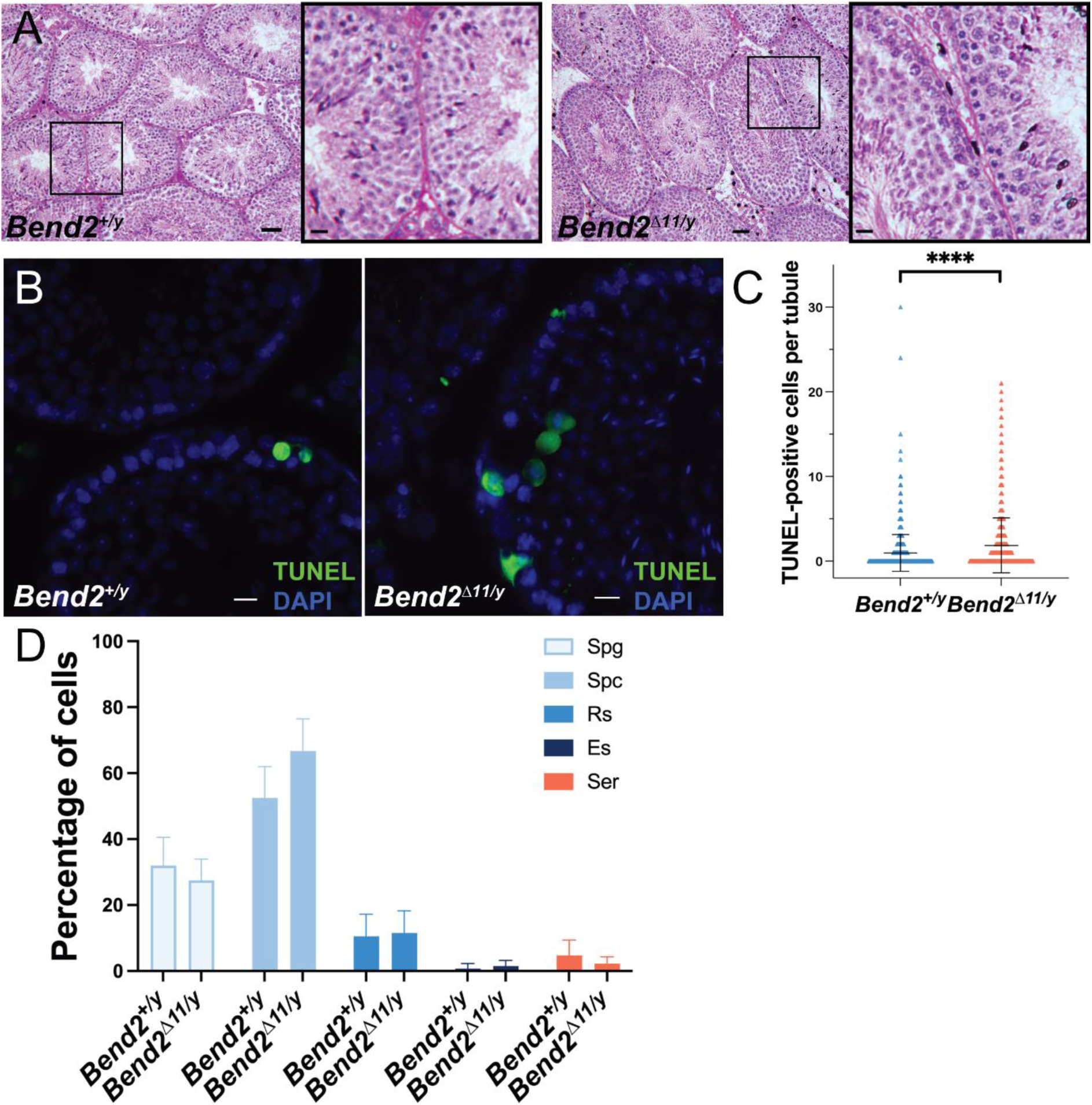
The full-length BEND2 is dispensable to complete spermatogenesis in mice. (A) Representative PAS-H stained mouse testis sections. The square on the left image shows the zoomed images on the right. Scale bar on the right images, 20 µm. (B) Apoptosis detection on testis sections by TUNEL assay. Scale bar, 20 µm. (C) Quantification of TUNEL-positive cells. The horizontal lines represent the mean ± SD. N=1125 for *Bend2^+/y^*; N=1195 for *Bend2^Δ11/y^*, \*\*\*\**p*<0.0001 t-test. (D) Classification of TUNEL-positive cells. The columns and error lines indicate the mean ± SD. N=4, *p*>0.05 One-Way ANOVA.

To further explore the causes of the increased apoptosis in BEND2–deficient testes, we assessed chromosome synapsis and meiotic recombination progress in spermatocytes by IF. In wild-type spermatocytes, SYCP3 began to form each developing chromosome axis at leptotene. At zygotene, synapsis initiated as SYCP1 appeared at the synapsed region of the homologs. At pachytene, synapsis was completed as SYCP3 and SYCP1 completely colocalized. At diplotene, SCs disassembled, and SYCP1 were lost from separated SYCP3-labelled axes, but homologous chromosomes remained held together by chiasmata.

Synapsis progressed similarly to wild-type spermatocytes in *Bend2^Δ11/y^* spermatocytes (**Fig 4A**). However, the fraction of diplotene spermatocytes was significantly increased (*p*=0.019, One-Way ANOVA). Additionally, the fraction of pachytene spermatocytes (48.6 ± 6.8%, mean ± SD, N=4) seemed to decrease in *Bend2^Δ11/y^* mice, compared to wild-type mice (57.3 ± 4.5%, mean ± SD, N=4, *p*=0.078 One-Way ANOVA). This accumulation of *Bend2^Δ11/y^* spermatocytes at the diplotene stage could be explained if there was a later block of the meiotic progression or if *Bend2^Δ11/y^* spermatocytes might exit pachytene faster than wild-type spermatocytes. Taken together, these results demonstrated that in the absence of the full-length BEND2, spermaocytes could complete synapsis and progress through meiotic prophase but following a slightly altered timeline.

**Figure 4.**
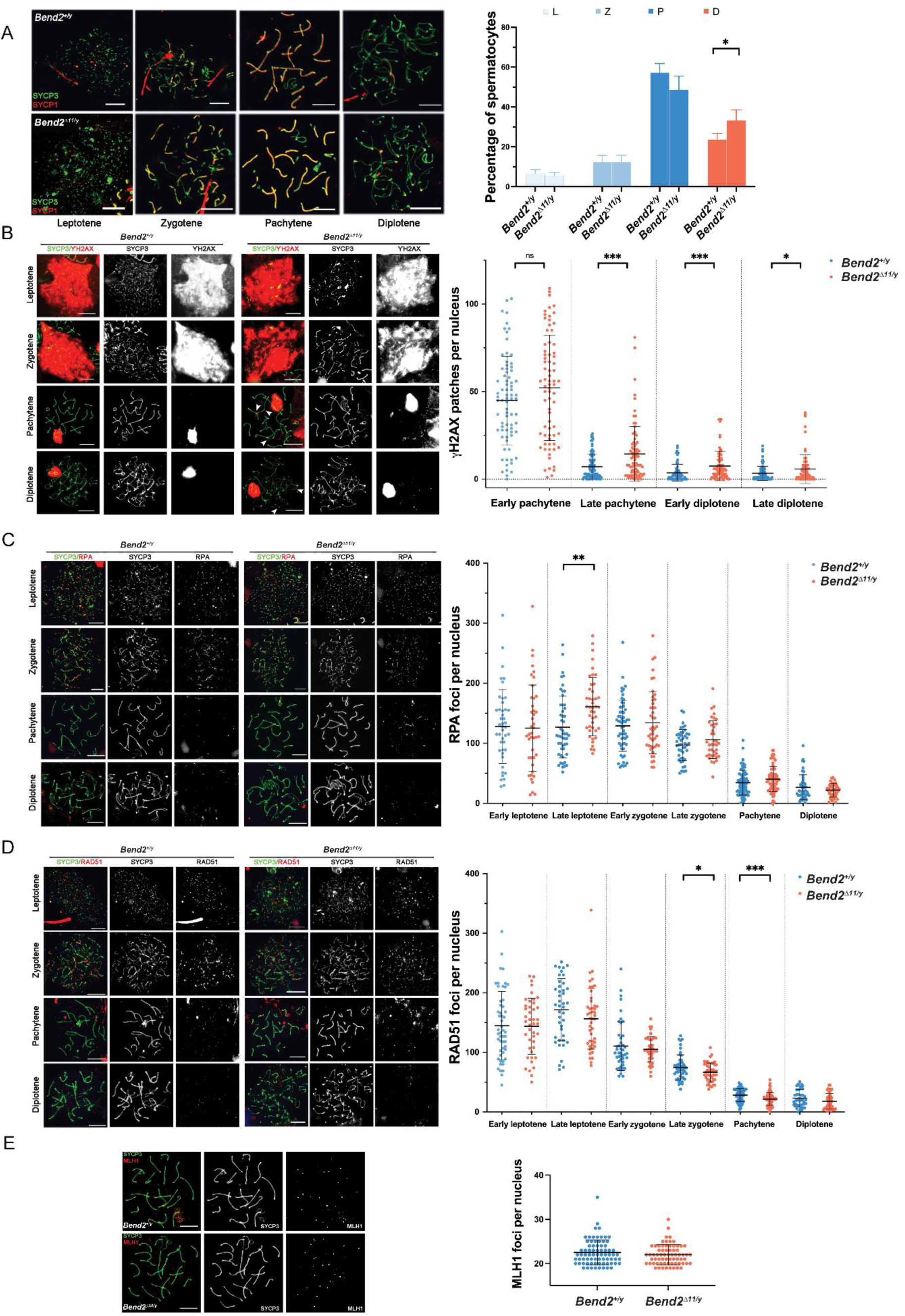
*Bend2^Δ11/y^* males display minor recombination defects. (A) Chromosomal synapsis in spermatocytes. Representative images of SYCP3 and SYCP1 staining in spermatocyte nuclei from the stages shown (left). Meiotic prophase staging of spermatocytes (right). L: leptotene, Z: zygotene, P: pachytene, D: diplotene. The columns and lines indicate the mean and SD. N=4, \**p*=0.019 One-Way ANOVA. (B) Examination of DSBs in *Bend2^Δ11/y^* spermatocytes. Representative images of ϒH2AX staining in spermatocyte nuclei along meiotic prophase (left). Increased ϒH2AX signals are detected during late prophase in *Bend2^Δ11/y^* spermatocyte (white arrowhead). Quantification of ϒH2AX patches in spermatocyte nuclei per sub-stages (right). The patches were counted manually using the same method for every pair of control and mutant mice. From left to right, N=76/74, 86/83, 73/75, and 78/81, \*\*\**p*=0.0001 (late pachytene) and 0.0007 (early diplotene), \**p*=0.0222 t-test. Analysis of RPA (C) and RAD51(D) foci counts in control and *Bend2^Δ11/y^* spermatocytes. RPA and RAD51foci were counted using ImageJ with the same method for every *Bend2^+/y^* and *Bend2^Δ11/y^* mice pair. From left to right, N=44/43, 51/43, 59/46, 45/39, 81/77, and 48/43, \*\**p*=0.0015 for RPA quantification; N=52/43, 42/46, 44/46, 51/45, 77/73, and 45/49, \**p*=0.0428 and \*\*\**p*=0.0002 for RAD51 quantification; t-test. C) Examination of CO formation in *Bend2^Δ11/y^* spermatocytes. Representative images of MLH1 in control and mutant spermatocyte nuclei (left). Quantification of MLH1 foci in spermatocyte nuclei (right). Only spermatocytes containing ≥ 19 MLH1 foci/nucleus were counted. N=74 for *Bend2^+/y^* and 68 for *Bend2^Δ11/y^*, *p*>0.05 Mann-Whitney test. The horizontal lines represent mean ± SD (B-E). Scale bar, 10 µm.

During early meiosis, recombination initiates with SPO11-mediated DSB formation, leading to the phosphorylation of the histone H2AX by ATM and thus triggering a series of DSB repair responses in the meiotic prophase (Huang and Roig, 2023). In wild-type mice, ϒH2AX progressively disappeared from the autosomes while DSBs were repaired as prophase progressed. By pachytene, most ϒH2AX was associated with the sex body. Few ϒH2AX patches could be observed on the autosomes from late pachytene, presumably corresponding to unrepaired DSBs (**Fig 4B**).

Interestingly, a marked increase in the number of ϒH2AX patches was detected from early pachytene until late diplotene in *Bend2^Δ11/y^* spermatocytes, indicating a possible delay in DSB repair (**Fig 4B**). Alternatively, the same results could be observed if more DSBs were formed during prophase in *Bend2^Δ11/y^* spermatocytes.

Once DSBs are resected, replication protein A (RPA) transiently binds to the nascent ssDNA overhangs, promoting the assembly of the recombinases RAD51 and DMC1 at DSB sites (Moens et al., 2002). RAD51 and DMC1 replace RPA and form nucleoprotein filaments with the ssDNA, directing homology search and strand invasion to proceed along the DSB repair pathway (Brown and Bishop, 2015; Hinch et al., 2020).

In wild-type spermatocytes, the number of RPA foci peaked around early zygotene and progressively diminished as recombination proceeded; on the other hand, there were many RAD51 foci since leptotene, and the number started to decline from early zygotene (**Fig 4C-D**), consistent with previous studies (Moens et al., 2002; Pacheco et al., 2015). In *Bend2^Δ11/y^* males, the spermatocytes had similar initial RPA and RAD51 foci numbers, suggesting that DSB formation was not affected by the loss of BEND2 (**Fig 4C-D**). However, *Bend2^Δ11/y^* cells tended to accumulate more RPA foci and lose RAD51 foci as they progressed along the meiotic prophase. The differences became significant for RPA at late leptotene (*p*=0.0015 t-test, **Fig 4C**) and RAD51 at late zygotene and pachytene (*p*=0.0428 and 0.0002 respectively t-test, **Fig 4D**), indicating a subtle defect in the replacement of RPA by RAD51.

Because of meiotic recombination, at least one crossover per bivalent is generated to ensure accurate chromosome segregation at the first meiotic division (Zickler and Kleckner, 1999). We examined MLH1 foci, which become apparent at mid-late pachytene and mark most crossover-designated sites (Anderson et al., 1999). We did not detect significant differences in the number of MLH1 foci in *Bend2^Δ11/y^* spermatocytes (22.0 ± 2.3, mean ± SD, N=68) compared to wild-type cells (22.5 ± 2.9, mean ± SD, N=74, *p*=0.2543, t-test, **Fig 4E**), suggesting that BEND2 deficiency did not affect crossover formation in spermatocytes.

Homologous recombination is the principal DSB repair pathway in meiosis responding to SPO11-dependent DSBs. But in late spermatocytes (late pachytene and diplotene), other DSB repair pathways, such as non-homologous end joining (NHEJ) or inter-sister homologous recombination (IS-HR), take over (Ahmed et al., 2010; Enguita-Marruedo et al., 2019; Goedecke et al., 1999). Moreover, during pachytene, new DSBs could be formed by SPO11-independent mechanisms (Carofiglio et al. 2013). Therefore, the increased presence of ϒH2AX patches observed in *Bend2* mutant mice might be explained by the inactivation of these alternative DSB repair pathways during the late meiotic prophase or an increase of SPO11-independent DSBs during pachytene.

To gain insight into these hypotheses, we studied if the NHEJ pathway activity is reduced during meiosis in *Bend2^Δ11/y^* mice by examining the localization and expression of its constitutive protein, Ku70, on testes sections. As anticipated, Ku70 was observed as a patch-like signal on sex bodies in pachytene and diplotene spermatocytes. A punctuated signal, marking presumably DSB sites in the nucleus, was seen in cells of all spermatogenesis stages besides round and elongated spermatids (**Fig S2A**). The number of Ku70 foci colocalizing with SYCP3 in spermatocytes in pachytene and diplotene stages was similar between control and mutant samples (**Fig S2B**). No differences were observed in the proportion of tubules expressing Ku70 either. These results were further confirmed with Western blot analysis (**Fig S2C and D**), thus suggesting that *Bend2* depletion does not affect NHEJ activation.

### BEND2 deficiency leads to LINE-1 retrotransposon suppression

The formation of SPO11-independent DSBs during meiotic prophase has been linked to the LINE-1 retrotransposon activation (Carofiglio et al., 2013; Malki et al., 2014; Soper et al., 2008). To evaluate if the LINE-1 activity was increased in *Bend2* mutant mice, we examined LINE1 expression in mouse testes by IF and WB.

As expected, the most prominent LINE-1 staining localized to the cytoplasm of leptotene, zygotene, and pachytene spermatocytes (**Fig S3A**). The signal intensity varied greatly between seminiferous tubules of the same sample in both wild-type and mutant animals. In *Bend2^Δ11/y^* mice, spermatocytes showed similar signal intensity compared to control mice. Contrary to the expected, a statistically significant lower number of LINE-1 positive tubules was observed in mutant mice. An average of 2.8% and 0.5% positive tubules was observed in control and *Bend2^Δ11/y^* mice, respectively (*p*=0.02, t-test) (**Fig S3B**). Western blot analysis validated the lower LINE-1 protein expression in *Bend2^Δ11/y^* testes sections (**Fig S2B**). Indeed, a significant decrease in LINE-1 protein levels was found in *Bend2^Δ11/y^* mice (*p*=0.0469, t-test, **Fig S3C**), thus confirming the IF results. Defects in the demethylation efficiency of LINE-1 in *Bend2* mutant mice during the prophase may explain these results. On the other hand, the signal intensity was equal in wild-type and mutant mice, suggesting that some tubules successfully complete LINE-1 demethylation, thus resulting in normal LINE-1 expression.

### BEND2 deficiency results in a reduced ovarian reserve

A more robust phenotype was observed in females when BEND2 was disrupted compared to males. Female *Bend2^Δ11/Δ11^* mice were fertile. However, the litter size was significantly smaller than that in wild-type females (**Fig 5A**), and noticeably smaller ovaries were present in mutant females (**Fig 5B**).

**Figure 5.**
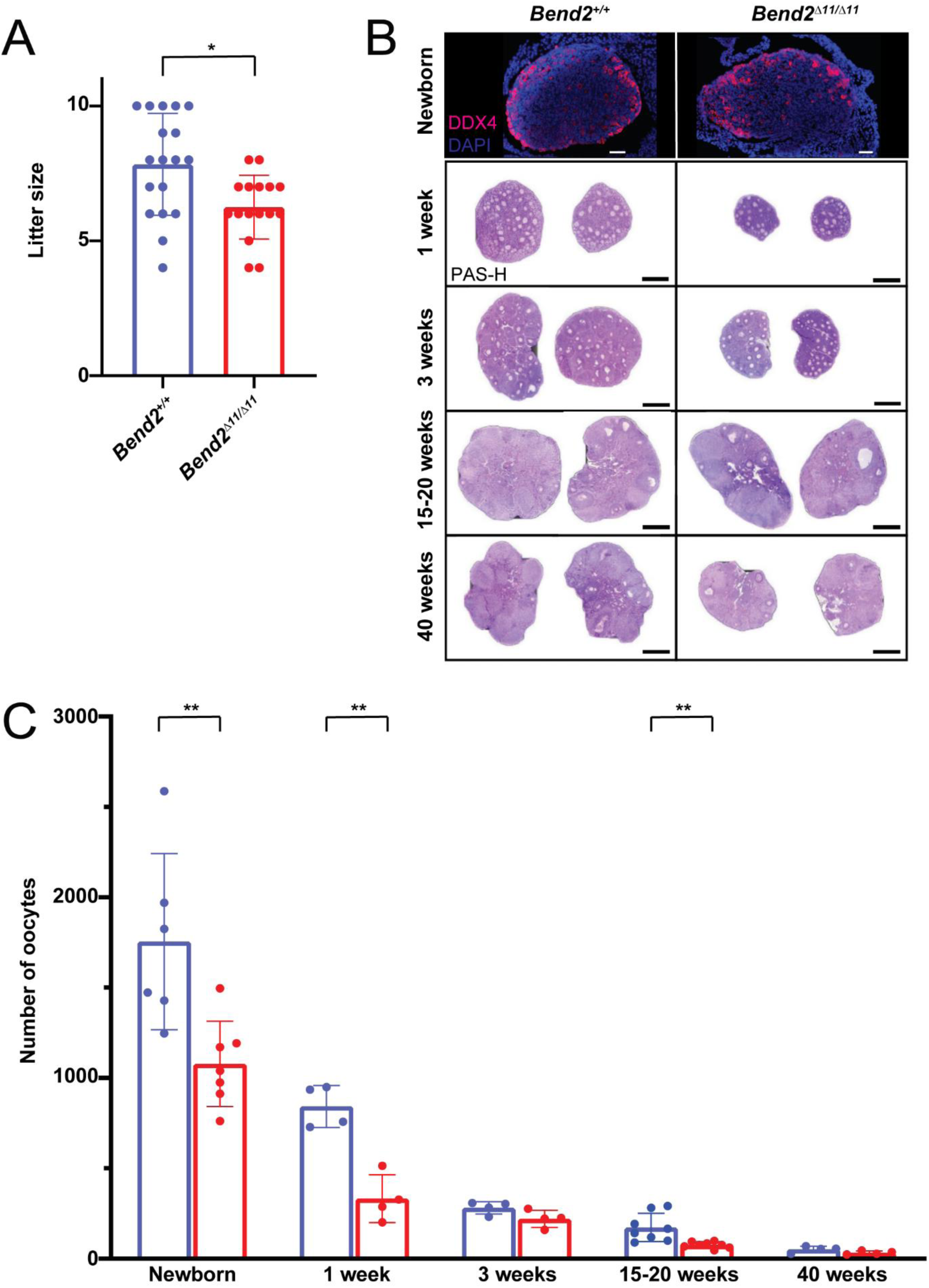
Oogenesis is altered in *Bend2* mutant females. (A) Fertility evaluation of *Bend2^Δ11/Δ11^* females. Two-month-old *Bend2^Δ11/Δ11^* females were crossed with wild-type males for 5 months; litter size data of 4 animals per genotype were collected for analysis. \**p*=0.0069, t-test. (B) DDX4-stained and PAS-H stained histological ovary sections from females at different ages. Scale bars, 50µm for Immunofluorescent images and 0.5 mm for the PAS-H stained ones. (C) Quantification of the number of ocoytes found in the analysed ovaries of newborn, one, three, 15-20 and 40 weeks old mice. The columns and lines indicate the mean and SD. \*\**p*=0.0075, 0.0012 and 0.0040; t-test.

To investigate if the loss of BEND2 affected oogenesis, a more detailed histological analysis of whole ovaries was performed in young (1 week old), adolescent (3 weeks old), adult (15-20 weeks old), and aged (40 weeks old) *Bend2^Δ11/Δ11^* and *Bend2^+/+^* mice. At 1 week, *Bend2^Δ11/Δ11^* females exhibited a significantly reduced number of oocytes (*p*=0.0012, t-test; **Fig 5C**) mainly due to a reduced number of primordial follicles (*p*=0.0007, t-test; **Fig S4A**). Growing follicles were found at all stages, with no statistically significant differences between *Bend2^Δ11/Δ11^* and control mice (**Fig S4A**). This tendency continued in prepubertal 3-week-old animals. At this stage, Bend2 mutant ovaries seemed to contain fewer oocytes (*p*=0.0834, t-test; **Fig 5C**) and primordial follicles (*p*=0.0704, t-test; **Fig S4B**). At an adult stage of 15-20 weeks, the mutant ovaries presented significantly fewer oocytes (*p*=0.0040, t-test; **Fig 5C**), again due to a reduction in the number of primordial follicles (p=0.0024, t-test; **Fig S4C**). Interestingly, at this stage, we also noticed a significant reduction in the number of antral follicles in mutant mice (*p*<0.0001, t-test; **Fig S4C**), which could partly explain the reduced fertility of these females. Finally, at 40 weeks, there tended to be fewer oocytes and primordial follicles in *Bend2^Δ11/Δ11^* ovaries than in *Bend2^+/+^* ovaries (*p*=0.1171 and p=0.1383, t-test; **Fig S4D**). These results suggest a reduction in the oocyte pool established at birth and the possible occurrence of premature ovarian insufficiency in *Bend2* mutant females.

To certify that BEND2 was required to establish the ovarian reserve, we counted the oocyte population present in newborn *Bend2^+/+^* and *Bend2^Δ11/Δ11^* mutant mice by staining ovarian sections with the germ-cell marker DDX4 (**Fig 5B**). *Bend2* mutant ovaries contained almost 40% fewer oocytes than control ovaries (*p*=0.0037, t-test ; **Fig 5C**), evidencing that the absence of the full-length BEND2 resulted in a reduced ovarian reserve.

### Mutant *Bend2* oocytes show persistent unrepaired DSBs and impaired crossover formation

To address the cause of this reduced ovarian reserve, we first analyze the expression of LINE-1, which has been associated with increased perinatal oocyte death in mice (Malki et al., 2019). However, we found similar percentage of oocytes expressing LINE-1 in *Bend2^+/+^* and *Bend2^Δ11/Δ11^* ovarian sections (20.2 ± 0.6; 16.8 ± 2.3, mean ± SEM; p=0.1851, t-test ; **Fig S5**). These data suggest that the reduction in the number of oocytes present in *Bend2* mutant mice might not be related to an increased expression of LINE-1.

Next, we examined if errors during the meiotic prophase could be responsible for the observed reduced ovarian reserve in *Bend2^Δ11/Δ11^* mice. Thus, we studied chromosome synapsis and meiotic recombination progress in fetal oocytes by IF. In wild-type ovaries, most oocytes at 18 dpc were at the pachytene stage, a small fraction was still at the zygotene stage, and a minority at the leptotene stage. At 1 dpp, almost all oocytes were at the pachytene and diplotene stages (**Fig 6A**). In *Bend2^Δ11/Δ11^* ovaries, zygotene, and pachytene oocytes undergoing normal synapsis and diplotene oocytes undergoing desynapsis were observed at 18 dpc and 1 dpp (**Fig 6A**).

**Figure 6.**
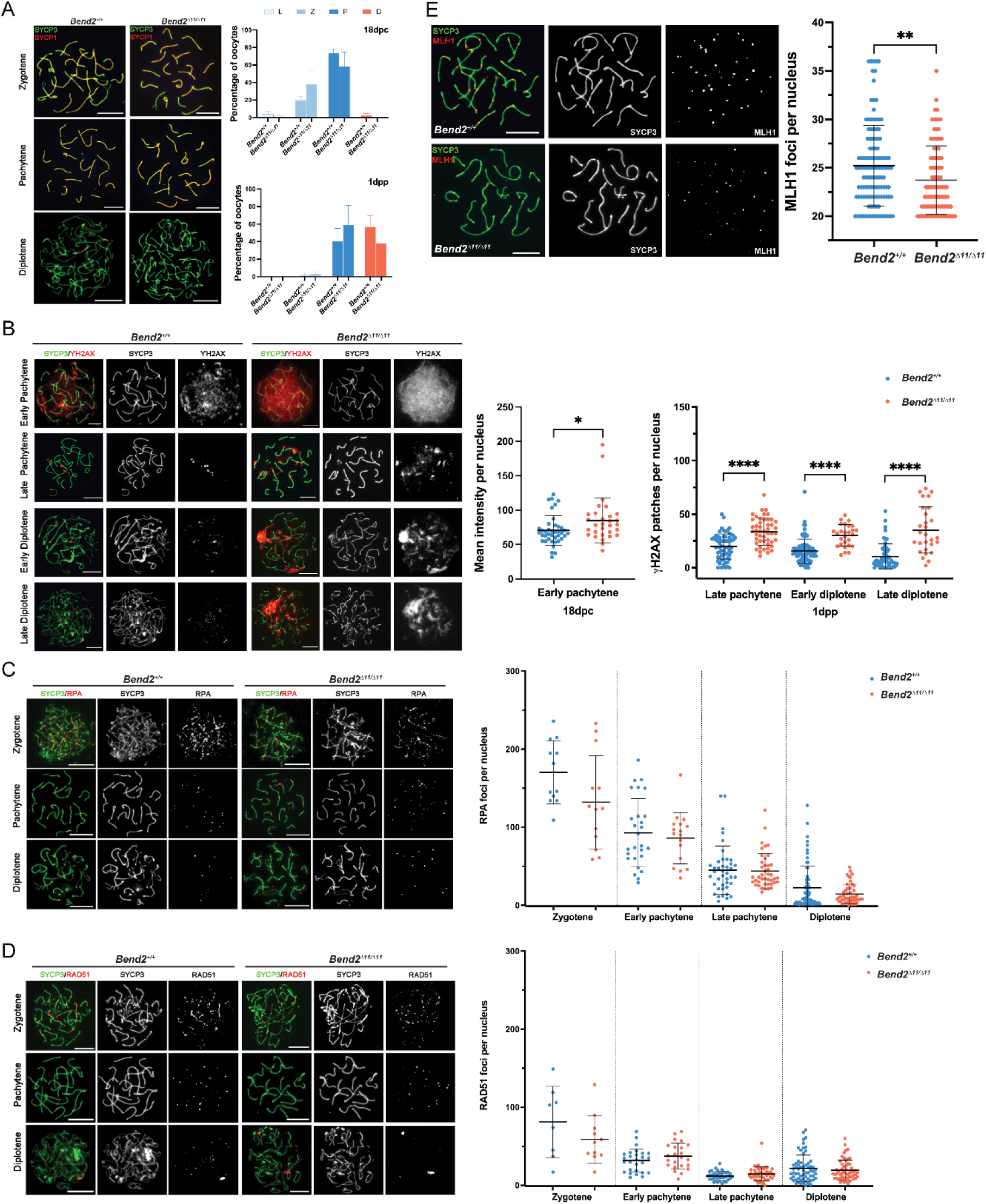
Synapsis and recombination in *Bend2^Δ11/Δ11^* females. (A) Chromosomal synapsis in oocytes. Representative images of SYCP3 and SYCP1 staining in 18 dpc and 1 dpp oocyte nuclei are shown (left). Meiotic prophase staging of 18 dpc and 1dpp oocytes (right). L: leptotene, Z: zygotene, P: pachytene, D: diplotene. The columns and lines indicate the mean and SD. The number of animals analyzed per genotype, 2 for 18 dpc and 3 for 1 dpp. *P*>0.5 for all the comparisons One-Way ANOVA. (B) Examination of DSBs in *Bend2^Δ11/Δ11^* oocytes. Representative images of ϒH2AX staining in 18 dpc and 1 dpp oocyte nuclei at the pachytene and diplotene stage (left). Quantification of ϒH2AX patches in oocyte nuclei at sub-stages (right). The mean intensity of ϒH2AX staining in 18 dpc oocyte nuclei was measured by Image J. The patches of ϒH2AX in 1 dpp oocyte nuclei were counted manually. The horizontal lines represent the mean ± SD. From left to right, the number of analyzed nuclei per genotype: 40/30, 64/48, 81/27, and 57/28. \**p*=0.0288, \*\*\*\**p*<0.0001 t-test. (C) Analysis of RPA (C) and RAD5 (D) foci in control and *Bend2^Δ11/Δ11^* oocytes. Representative images of RPA staining in 18 dpc and 1 dpp oocyte nuclei from zygotene to diplotene stage (left). Quantification of RPA foci present in oocyte nuclei at sub-stages (right). Zygotene and early pachytene nuclei analyzed were from 18 dpc ovaries, and late pachytene and diplotene nuclei analyzed were from 1 dpp ovaries. Foci were counted manually. The horizontal lines represent the mean ± SD. From left to right, number of analyzed nuclei: 12/13, 27/17, 41/42, and 65/49 for RPA, and 8/11, 27/23, 38/46, and 62/55 for RAD51. *p*>0.5 for all the comparisons t-test. (E) Examination of CO formation in control and *Bend2^Δ11/Δ11^* oocytes. Representative images of MLH1 in 1 dpp oocyte nuclei (left). Quantification of MLH1 foci in 1 dpp oocyte nuclei (right). Only oocytes containing ≥20 MLH1 foci/nucleus were counted. The horizontal lines represent the mean ± SD. N=120 for *Bend2^+/+^*, 95 for *Bend2^Δ11/Δ11^*; \*\**p*=0.0057 Mann-Whitney test. Scale bar, 10 µm.

Consistent with the observation in *Bend2* mutant males, we also found increased levels of ϒH2AX in late prophase *Bend2^Δ11/Δ11^* oocytes (**Fig 6B**). In wild-type females, ϒH2AX patches could be detected along axes in most pachytene oocytes and many diplotene oocytes (**Fig 6B**). This is different from wild-type males, in which ϒH2AX can only be detected as very few patches in late pachytene cells and is almost undetectable in most diplotene cells (**Fig 4B**), presumably because synapsis progresses faster than recombination in females than in males (Roig et al., 2004).

In *Bend2^Δ11/Δ11^* oocytes, higher levels of ϒH2AX were observed at all stages analyzed (early pachytene, late pachytene, and diplotene, **Fig 6B**). Many early pachytene *Bend2^Δ11/Δ11^* oocytes exhibited an overall intense ϒH2AX signal reminiscent of wild-type zygotene cells (**Fig 6B**). To quantitatively compare these differences, we measured the ϒH2AX signal intensity in early pachytene oocytes and counted the ϒH2AX patches in late pachytene and diplotene oocytes. As expected, a significant increase of ϒH2AX was present in *Bend2^Δ11/Δ11^* oocytes at all the tested stages (*p*=0.0288 for early pachytene; *p*<0.0001 for late pachytene, early diplotene and late diplotene, t-test, **Fig 6B**). Thus, like in *Bend2* mutant males, DSB repair was impaired, or more DSBs were formed in *Bend2^Δ11/Δ11^* females.

In females, no differences in the number of RPA and RAD51 foci were found between wild-type and *Bend2^Δ11/Δ11^* oocytes, apart from a slight decrease in RPA foci in *Bend2^Δ11/Δ11^* oocytes (**Fig 6C**) and a slight increase of RAD51 foci in pachytene *Bend2^Δ11/Δ11^* oocytes (**Fig 6D**). These results suggest that the increased presence of ϒH2AX in *Bend2^Δ11/Δ11^* oocytes may not be related to defective meiotic recombination but to other causes, like meiotic silencing of unsynapsed chromosomes (MSUC) or alternative DNA repair pathways.

Interestingly, the number of MLH1 foci in late prophase oocytes was significantly lower in *Bend2^Δ11/Δ11^* females than in control females (*p*=0.006 for *Bend2^Δ11/Δ11^*, t-test, **Fig 6E**). Thus, we concluded that the depletion of BEND2 caused a reduced crossover formation in oocytes, probably partly causing the reduced litter size in *Bend2* mutant females.

Finally, to check if these alterations affected *Bend2^Δ11/Δ11^* oocyte quality, we performed ovarian stimulation on 7-month-old *Bend2^+/Δ11^* and *Bend2^Δ11/Δ11^* mice. On average, we obtained a similar number of oocytes from the ovarian stimulation in control and mutant mice (10.7 ± 4.1 vs. 10.0 ± 1.2; mean ± SEM; p=0.8852, t-test; **Table 1**). Remarkably, the ability of *Bend2^Δ11/Δ11^* mutant oocytes to be fertilized and develop into the blastocyst stage did not seem to be compromised (**Table 1**), suggesting that the absence of the full-length BEND2 did not compromise oocyte quality, at least to be fertilized and develop up to the blastocyst stage.

**Table 1.** *Bend2^Δ11/Δ11^* oocytes fertilize and develop into blastocyst stage similarly than control oocytes.

| Group | Genotype | # mouse | Age (months) | # oocyte | # 2-cell embryo | Fertilization (%) | Blastocyst (%) |
| --- | --- | --- | --- | --- | --- | --- | --- |
| Control | *Bend2*<sup>+/$\Delta 11$</sup> | 3 | 7 | 10.7±7.2 | 7.7±3.8 | 78.0±19.5 | 23.3±25.2 |
| Mutant | *Bend2*<sup>$\Delta 11/\Delta 11$</sup> | 3 | 7 | 10.0±2.0 | 8.3±3.2 | 82.7±20.5 | 46.8±12.2 |

## Discussion

In mammals, the establishment and gradual depletion of the ovarian reserve defines a finite female reproductive life span. In this study, we demonstrated that BEND2, a new player in meiosis (Ma et al., 2022), is essential for the primordial follicle pool setup. The depletion of the full-length BEND2 causes severe defects in female oogenesis with a decreased ovarian reserve and subfertility. However, these defects may not be directly related to meiotic recombination progression or synapsis, although a high level of unrepaired DSBs persists during late meiotic prophase. We also showed that BEND2 has a role in regulating LINE-1 retrotransposon activity in spermatocytes. Our work expands our knowledge of the genetic factors influencing the ovarian reserve, potentially benefiting the genetic diagnosis of human infertility.

BEND2 has two C-terminal BEN domains, marked by α-helical structure, and is conserved in a range of metazoan and viral proteins. BEN domain is suggested to mediate protein–DNA and protein-protein interactions during chromatin organization and transcription (Abhiman et al., 2008). So far, only a few BEN domain-containing proteins have been described. All of their functions are linked to transcriptional repression regardless of whether the proteins contain other characterized domains, including mammalian BANP/SMAR1, NAC1, BEND3 and RBB and *Drosophila* mod(mdg4), BEND1 and BEND5 (Dai et al., 2015, 2013; Gerasimova et al., 1995; Kaul-Ghanekar et al., 2004; Korutla et al., 2007, 2005; Nègre et al., 2010; Rampalli et al., 2005; Sathyan et al., 2011; Xuan et al., 2012). BEND1, BEND5, and RBB are revealed to bind DNA through sequence-specific recognition (Dai et al., 2015, 2013; Xuan et al., 2012), and BEND3 specifically binds heterochromatin (Sathyan et al., 2011a). Additionally, several human BEND2 fusion proteins have been identified in tumors, and the activation of BEND2 is suggested to promote oncogenic activity (Burford et al., 2018; Scarpa et al., 2017; Sturm et al., 2016; Williamson et al., 2019). These data suggest that BEND2 might have a role in regulating chromatin and transcriptional gene expression. A recent study showed that BEND2 is a crucial regulator of meiosis gene expression and chromatin state during mouse spermatogenesis (Ma et al., 2022).

In the study of Ma et al. (2022), two BEND2-related proteins (p140 and p80) were found in mouse testis, using an antibody against a short polypeptide from the C-terminus of BEND2. They demonstrated that p140 is the full-length version of BEND2, displaying slower electrophoretic mobility due to its unusual sequence/structure at the N-terminus. They also reported that BEND2 was expressed in the nuclei of spermatogenic cells around meiosis initiation. Similarly, using our antibody against the whole C-terminus of the protein, we also found two BEND2-related proteins expressed in mouse testis and a highly consistent expression pattern of BEND2 during spermatogenesis. These two proteins likely correspond to full-length and smaller isoform from Ma et al. (2022).

Ma et al. (2022) disrupted *Bend2* by deleting exons 12 and 13, which resulted in the loss of both the full-length (p140) and the smaller isoform (p80) of BEND2. In their study, *Bend2* mutant mice were sterile and exhibited severe meiotic recombination and synapsis defects. In contrast, our study targeted exon 11, specifically ablating the full-length p140 isoform while preserving the expression of the smaller p80 isoform in the testis. Interestingly, *Bend2^Δ11/y^* mutant males remain fertile, with essentially normal meiotic recombination and synapsis, suggesting that the full-length BEND2 (p140) is not essential for the repair of SPO11-induced double-strand breaks or for synapsis during meiosis.

This observation strongly implies that p80 is sufficient to fulfill the essential roles of BEND2 in meiosis and spermatogenesis. The preserved fertility and proper progression of meiosis in our *Bend2^Δ11/y^* mutants suggest that the smaller p80 isoform is the primary driver of BEND2’s function in these processes. Conversely, the sterility and severe meiotic defects observed in the study by Ma et al. can be attributed to the loss of p80 and p140. These findings highlight a functional redundancy or compensatory mechanism provided by p80 that allows meiosis and spermatogenesis to proceed effectively in the absence of the full-length isoform.

Thus, we propose that p80 is the indispensable isoform of BEND2 required for meiotic progression. Its presence in our model likely prevents the spermatogenesis arrest seen in the Ma et al. study. This hypothesis provides a compelling explanation for why our *Bend2^Δ11/y^* males exhibit a less severe phenotype and remain fertile despite the absence of full-length BEND2. These results underscore the importance of considering isoform-specific functions when investigating the roles of multi-isoform genes in complex biological processes like meiosis and gametogenesis.

In our study, one common defect in mutant spermatocytes and oocytes was a significant increase in unrepaired DSBs during late prophase. Examination of the chromosomal dynamics and meiotic recombination demonstrated that the full-length BEND2 is not essential for SPO11-induced DSB repair. Since all meiotic recombination markers studied, apart from γH2AX, were mostly indistinguishable in mutant and control cells, we hypothesized that the observed unrepaired DSBs might be SPO11-independent and further explored the LINE-1 retrotransposon activation, which is known to cause DSBs during meiosis (Soper et al., 2008; Carofiglio et al., 2013; Malki et al., 2014) and it has been implicated in the regulation of the ovarian reserve (Martínez-Marchal et al., 2020; Tharp et al., 2020). Unexpectedly, we found a significant reduction in LINE-1 expression levels in mutant testis and although it was not statistically significant, mutant oocytes seemed to have a similar trend.

Repetitive transposon elements, including LINE-1, occupy almost half of the human genome and are extremely important for genome evolution (Wang, 2017). Transposon-mediated insertional mutagenesis was linked to a multitude of genetic diseases. Even though LINE-1 is active mainly during the genome-wide demethylation, transient DNA demethylation occurring at the onset of meiosis was found to trigger a persisting LINE-1 expression until mid-pachytene (Soper et al., 2008). Silencing of LINE-1 retrotransposons is achieved with multiple mechanisms such as DNA methylation, histone modifications, and piRNAs (piwi-interacting RNAs) (DiGiacomo et al., 2013). Ma et al. (2022) described that in *Bend2* KO male mice, there is an up-regulation of PIWIL2 expression at the onset of meiosis. PIWIL2 belongs to the PIWI subfamily of proteins that bind to piRNAs, which are required for several biological processes, including silencing retrotransposons during meiosis (DiGiacomo et al., 2013). Thus, it is likely that *Bend2* depletion caused an up-regulation of the piRNA pathway, resulting in lower LINE-1 expression in our mouse mutant spermatocytes. Alternatively, as BEND2 was found to bind to repetitive sequences (Ma et al., 2022), it could interact directly with LINE-1 to promote its expression. The lower LINE-1 expression might result straightforwardly from less binding of BEND2. Finally, *Bend2* mutation could affect chromatin state and higher-order structures (Ma et al., 2022), which could alter LINE-1 expression and hinder DNA repair. Further studies focusing on the regulation of PIWIL2 by BEND2 will clarify these aspects of the phenotype.

Due to the sterility of hemizygous mutant males, female homozygous mutants can’t be produced in the study of Ma et al. (2022). So, our female mutant model facilitates further exploration of BEND2’s roles in oogenesis.

Interestingly, our mutant females *Bend2^Δ11/Δ11^* exhibited a more severe phenotype with a reduced ovarian reserve and subfertility, which had not been described before. Mouse females establish the pool of primordial follicles around birth. As folliculogenesis begins, the pool is gradually depleted throughout reproductive life (Hunter, 2017). By examining the dynamics of the oocyte pool in female mice at different ages, we showed that compared to wild-type females, BEND2-deficient females have a significantly smaller ovarian reserve. As a result, fewer follicles develop during folliculogenesis from pre-puberty throughout adulthood. Consistent with these, BEND2-deficient females produced reduced litter size, which indicates subfertility. Interestingly, our mutant oocyte quality analysis suggests that mature oocytes from mutant females are equally competent to develop into a blastocyst as control ones. These data suggest that the subfertility observed in *Bend2* mutants may be due to errors in later developmental stages, such as implantation or organogenesis.

Thus, BEND2, apart from its roles in male meiosis (Ma et al., 2022), is also required to establish the ovarian reserve during oogenesis. The full-length BEND2 may take on this role in females. However, future investigation of BEND2 expression in fetal ovaries is necessary to validate this. Genes discovered to have disease-causing roles in mouse models often offer a panel of strong candidate genes for screening human infertility factors (Huang and Roig, 2023; Riera-Escamilla et al., 2019). Our data shows that the depletion of BEND2 may lead to premature ovarian insufficiency (POI), which is also a significant cause of female infertility in humans, and its underlying genetic causes are mainly unknown (Rossetti et al., 2017). Thus, identifying BEND2’s requirement for normal oogenesis is also important to recognizing the genetic determinants of human POI.

## Acknowledgments

This work was supported by Spanish Ministerio de Ciencia, Innovación e Universidades grants (I.R: PID 2019-107082RB-I00, PID2022-138905OB-I00; A.M.P.: PID2020–120326RB-I00), Junta de Castilla y León grants (A.M.P.: CSI017P23 and CSI017P23) and a grant from La Fundació Marató de TV3 (I.R.: 677/U/2021). Yan Huang is a recipient of a fellowship from the China Scholarship Council (201607040048). Cristina Madrid-Sandín is the recipient of an FPU fellowship from the Spanish Ministerio de Ciencia, Innovación e Universidades (FPU19/02885). Nikoleta Nikou is supported by an FPI fellowship from the Spanish Ministerio de Ciencia, Innovación e Universidades (PRE2020-094355). Moreover, we would like to acknowledge the rest of the members of the Roig Lab for their support, comments, and discussions about the project.

## Methods and materials

### Mice

*Bend2* mutant mice were generated by using the CRISPR/Cas9 system. A pair of gRNAs with minimum off-target and maximum on-target activity were designed and selected to specifically target sequences that encode essential protein domains of the *Bend2* gene using CRISPR DESIGN TOOLS (Millipore Sigma) (gRNA1-antisense: AGTAGCAGGCTGCATAAGT GGG; gRNA2-sense: AGACCAGCCTTATTGACCATGG). SgRNAs were synthesized by Sigma and microinjected with Cas9 protein into the pronucleus of C57BL/6JOlaHsd zygotes. Edited founders were identified by PCR with primers flanking the targeted region and HincII digest. PCR products were further purified and determined by Sanger sequencing. Five out of 17 F0 mice were identified to carry desired mutations, including two homozygous males and three heterozygous females. Each of them was crossed with wild-type C57BL/6JOlaHsd to eliminate possible off-target mutations to generate pure heterozygotes. Female heterozygotes (*Bend2^+/Δ11^*) were crossed with wild-type males (*Bend2^+/y^*) to obtain wild-type females (*Bend2^+/+^*), heterozygous females (*Bend2^+/Δ11^*), wild-type male (*Bend2^+/y^*), and mutant males (*Bend2^Δ11/y^*) offsprings. Female heterozygotes (*Bend2^+/Δ11^*) were crossed with mutant males (*Bend2^Δ11/y^*) to obtain mutant females (*Bend2^Δ11/Δ11^*), heterozygous females (*Bend2^+/Δ11^*), wild-type males (*Bend2^+/y^*) and mutant males (*Bend2^+/y^*). Mice from at least F2 generation were analyzed for their phenotyping. All experiments used at least three animals from each genotype (unless mentioned in the text). Mutant males were compared with their wild-type littermates. Testes from 2 to 6 months old mice were collected and processed for adult mice. Female mutants were compared to wild-type mice from other litters of the same age and from animals of closely related parents.

For genotyping, genomic DNA was extracted from mouse tails by overnight incubation at 56℃ in lysis buffer (0.1 M Tris-HCl pH 8.5-9, 0.2 M NaCl, 0.2% SDS, 5 mM EDTA and 0.4 mg/ml proteinase K), followed by precipitation with isopropanol and washes in cold 70% ethanol and subjected to PCR using NZYTaq II 2x Green Master Mix employing primer pair (forward: TTGCCAGTGGGGTATTACGA, and reverse: CTGGAAGGCAGGAAGTTTAACA). For female genotyping, the identified possible homozygous BEND2 females were subjected to an extra PCR using the primer pair (forward: TTTGCTCCACTGTTTCACGC, and reverse: TCCCTTTAAACTGCCAACAACA) to confirm the homozygosity.

The experiments performed in this study complied with EU regulations and were approved by the Ethics Committee of the UAB and the Catalan Government.

### Ovarian Stimulation and In Vitro Fertilization

Intra-peritoneal injection of 15IU pregnant marés serum gonadotrophin per female was administered to three *Bend2^+/Δ11^* and three *Bend2^Δ11/Δ11^* 7-month-old mice. After 49.5 h, 10 IU human chorionic gonadotrophin (hCG) per female injected and 13.5 hour later mice were sacrificed. Oviducts of treated females were dissected under stereo-microscope and cumulus masses were released into drop of fertilization medium (0.25mM GSH in HTF medium). Oocytes released from each female mice were counted. Time of ovulation was counted as 0.5dpc.

For in vitro fertilization, fresh sperm from a proven fertile, C57BL/6N wild-type male of 4 months of age were used and poured into dish containing oocytes in fertilization medium. After 3 hours of fertilization, oocytes were washed and O/N cultured in HTF medium. Assessment of fertilized, un-fertilized oocytes was done under stereo-microscope. From 2 cell-stage until blastocysts, embryos were cultured in KSOM medium.

### Total RNA purification and RT-PCR

Total RNA was purified from mouse tissues using the RNeasy Plus Mini Kit (Qiagen) and then transcribed into cDNA using the iScript™ cDNA Synthesis Kit (Bio-Rad), following the manufacturer’s instructions. cDNA was amplified using NZYTaq II 2x Green Master Mix with gene-specific primers (255P1 forward: TAGGGACCAAGAACCTGCTG, and reverse: TCCTGAAGCCACTGAGAAGG) and β-actin primers (forward: AGGTCTTTACGGATGTCAACG, and reverse: ATCTACGAGGGCTATGCTCTC) as control. A minus Reverse Transcription control (RT-) containing all the reaction components except the reverse transcriptase was included for testing for contaminating DNA.

### Gene cloning and transfection

cDNA from wild-type mouse testis was subject to PCR using Phusion High-Fidelity DNA Polymerase (Thermo Fisher Scientific) with a pair of specific primers (forward: agaaATGCCAGGAAAAACTGAAG, and reverse: TTAAGCTATTGCATTCCTTGGG for pEGFP-C1 vector and gAGCTATTGCATTCCTTGGGC for pEGFP-N1 vectors) targeting at both ends of the coding sequence to amplify the full length of *255p1(Bend2* novel splice variant). DNA purified from PCR reaction mix was phosphorylated and inserted into dephosphorylated pEGFP-C1 and pEGFP-N1 vectors. Plasmids were transformed to 5-alpha competent *E. coli* cells (NEB) and identified by colony PCR followed by Sanger sequencing.

pEGFP-C1-BEND2 or pEGFP-N1-BEND2 were transfected into the human embryonic kidney cell line (HEK 293T) by using jetPEI® DNA transfection reagent (Polyplus Transfection) and into 16-18 dpp wild-type live mouse testis by electroporation (Shibuya et al., 2014). Mice were sacrificed 24–72 hours post-electroporation and used for experiments.

### Antibody generation

To generate *in-house* BEND2 polyclonal antibody, the full length of 255p1(Bend2 novel splice variant) was amplified from pEGFP-C1-BEND2 and cloned into modified bacterial expression vector pET-28a (+)-TEV. The recombinant His-tagged BEND2 protein was expressed in *E. coli* BL21(DE3) competent cells by IPTG induction and then purified by affinity chromatography using HisTrap HP column (Cytiva). Purified His-tagged BEND2 protein was treated with TEV protease, followed by another affinity purification, to remove the His-tag. His-tag removed BEND2 protein was used to immunize rabbits. After immunization, the immune response was checked by ELISA. Antibodies were purified from rabbit serum by affinity chromatography using Affi-Prep protein A resin cartridge (Bio-Rad).

### Western blot

Total protein was extracted from mouse testis using RIPA lysis buffer (50 mM Tris pH 8, 1% Triton X-100, 0.1% SD, 150 mM NaCl, 1 mM EDTA, 0.5% Sodium Deoxycholate, 10 mM NaF, 1× Protease Inhibitor). Testis tissue was disrupted and homogenized thoroughly with a pestle in RIPA lysis buffer, followed by 10 min incubation at 95℃. Each lysate sample’s protein concentration was determined using Pierce™ BCA Protein Assay Kit (Thermo Scientific). 50 ug of total protein was loaded per well, and samples were separated by TGX-PAGE gel electrophoresis in Tris-Glycine-SDS buffer. Proteins were transferred to PVDF membranes (Bio-Rad) by wet electroblotting in Tris/glycine buffer. Membranes were blocked in 5% non-fat milk in PBS for 2 h at room temperature, followed by incubation of primary antibody diluted in blocking buffer overnight at 4°C. The next day, after three washes of PBST (PBS containing 0.05% Tween-20), membranes were incubated in anti-rabbit HPR conjugate antibody (1:5000, 170-6515, Bio-Rad) in PBS for 1 h at room temperature. ECL substrate (Bio-Rad) was used for chemiluminescent detection. Imaging was performed on ChemiDoc Touch Imaging system (Bio-Rad), and images were analyzed by Imagelab software (Bio-Rad). Primary antibodies used in WB: anti-BEND2 Rb (*in-house*, 3.2 μg/ml), anti-Ku70 Rb (abcam, 1:2000), anti-LINE1 ORF1p Rb (abcam, 1:2000), and anti-GAPDH Rb (abcam, 1:2000).

### Nuclei spreading and immunofluorescence

Spermatocyte nuclei spreading was prepared using frozen testes. Protocol was adapted from (Liebe et al., 2004): briefly, a small portion of testis was cut and minced thoroughly by a sterile blade in cold PBS (pH 7.4) containing 1 × protease inhibitor, PI (Roche Diagnostics) on a petri dish; cell mixture was transferred to a sterile Eppendorf and sat for 15 min to sediment; 25 µl of supernatant cell suspension was spread onto a glass slide and incubated with 80 µl of 1% Lipsol containing 1 × PI, for 15-20 min, followed by fixation in 150 µl of the PFA fixative solution (1% Paraformaldehyde pH 9.2-9.4, 15% Triton X-100, 1x PI) for 2 h at room temperature in a closed humid chamber. Oocyte nuclei spreading was prepared from fresh fetal and perinatal ovaries. Briefly, one pair of ovaries were dissected from each female under a stereomicroscope (Nikon SMZ-1); ovaries were first incubated in 500 µl of M2 medium (Sigma-Aldrich) containing 2.5 mg/ml collagenase (Sigma-Aldrich) for 30 min at 37℃, and then incubated in 500 µl of hypotonic buffer (30 mM Tris-HCl pH 8.2, 50 mM Sucrose, 17 mM Sodium Citrate, 5 mM EDTA, 0.5 mM DTT, 1x PI) for 30 min at room temperature; finally, single-cell suspension was prepared by disaggregating ovaries in 100 mM sucrose by pipetting under the stereo microscope. Each 10 µl of the cell suspension was spread onto a glass slide, followed by fixation in 40 µl of the PFA fixative solution (1% PFA, 5 mM Sodium Borate, 0.15% Triton X-100, 3 mM DTT, 1x PI, pH 9.2) for 2 h at room temperature in a closed humid chamber. Slides were dried under a fume hood and then washed in 0.4% Photoflo (Kodak) solution four times.

For immunofluorescence staining, slides of spermatocyte or oocyte nuclei spreading were blocked in freshly made blocking solution (0.2% BSA, 0.2% gelatin, 0.05% Tween-20 in PBS) at room temperature, followed by incubation of primary antibody diluted in blocking solution in a humid chamber overnight at 4℃; the next day, slides were washed four times in blocking solution, and incubated in secondary antibody diluted in blocking solution in a humid chamber for 1 h at 37℃, followed by another four washes in blocking solutions. Drained slides were mounted with 0.1 µg/ml DAPI in Vectashield antifade mounting medium. and analyzed with an epifluorescence microscope (Zeiss Axioskop). Primary antibodies used for IF: anti-SYCP3 Ms or Rb (abcam, 1:200), anti-SYCP1 Rb (abcam, 1:200), anti-phospho-Histone H2A.X Ms (Millipore, 1:400), anti-MLH1 Ms (BD Biosciences, 1:50), RPA32 (4E4) Rat (cell signaling, 1:100), anti-RAD51 (ab-1) Rb (Millipore, 1:100) and anti-GFP Rb (Thermo fisher scientific, 1:200).

### PAS-Hematoxylin, immunohistochemical and TUNEL staining

Fresh mouse tissues were fixed overnight at 4℃ with freshly made 4% PFA (0.4 g paraformaldehyde dissolved in 10 ml PBS, pH 7.4) for immunohistochemical (IHC) /TUNEL assay or with Bouin’s fixative for the use of PAS (Periodic Acid Schiff) staining. After fixation, tissues were washed in PBS, dehydrated in a series of ethanol with increasing concentration, cleared by histoclear and infiltrated by paraffin in sequence; embedded tissues were cut into thin slices (6-7 µm for testes; 4 µm for ovaries) using a microtome, which were mounted on poly-L-lysine coated slides and dry overnight at 37℃. Before any staining procedures, slides were deparaffinized in Xylene and a series of ethanol with decreasing concentration in sequence and rehydrated in distilled water.

For IHC, an antigen retrieval step was performed to expose the epitopes masked by PFA fixation: incubating the slides in sodium citrate buffer (10 mM Sodium citrate, 0.05% Tween 20, pH 6.0) or Tris-EDTA buffer (10 mM Tris, 1 mM EDTA, 0.05% Tween 20, pH 9.0) for 30 min at 95-100° C. Subsequently, an immunofluorescence was performed as described above and the used primary antibodies were anti-DDX4 (Abcam, 1:100) and anti-LINE-1 (Abcam, 1:100).

For TUNEL assay, tissue section samples were permeabilized in in 0.5% Triton X-100 in PBS for 15 min followed by two washes in PBS, incubated in background reducing solution (Dako) followed by a rinse in PBS, incubated in TUNEL reaction mixture (10% TdT enzyme solution, Roche Diagnostics) for 1 h at 37℃ in a humid chamber, followed by three washes in PBS. Slides were mounted with 15 µl DAPI (0.1 µg/ml in Vectashield antifade mounting medium) and analyzed with an epifluorescence microscope (Zeiss Axioskop).

For PAS-Hematoxylin (PAS-H) staining, tissue sections were oxidized by 1% Periodic Acid solution for 10 min, followed by two washes in distilled water, stained by Schiff’s reagent for 30 min in darkness, followed by two washes in sulfurous water (10% Potassium metabisulfite, 0.1 M HCl in Milli-Q water) and two washes in distilled water; counterstained in Mayer’s Hematoxylin for 1 min and rinsed in running tap water; dehydrated in a series of ethanal with increasing concentrations and xylene in sequence; mounted with DPX mounting medium and analyzed with an Optical microscope. Images were captured using Zeiss Axioskop microscope.

### Follicle count and classification

The whole ovaries were sectioned for follicle quantification in newborn, young, adolescent, and adult mice. Eight consecutive sections were mounted on each slide. Every third section was counted per slide to avoid counting follicles more than once. Five alternate slides per animal were counted (both ovaries). Follicles were only counted if the nucleus of the oocyte was visible. They were classified as primordial if they contained one single layer of plane squamous granulosa cells or as primary if they displayed one layer of cuboidal granulosa cells. Furthermore, follicles containing both plane and cuboidal cells were classified based on the predominant granulosa cell morphology. Follicles were classified as secondary if they showed more than one layer of cuboidal granulosa cells with no visible antrum. All follicles displaying an antral cavity of any size were classified as antral. Counting and classification were performed under a bright-field Zeiss Axioskop microscope.

### Image processing, analysis, and statistical analysis

All the images were processed by Adobe Photoshop. Fluorescence intensity was quantified by ImageJ; fluorescence signals were counted manually or by ImageJ. Data analysis and statistical inference were performed using GraphPad Prism 8 software.

---

**Figure S1.**
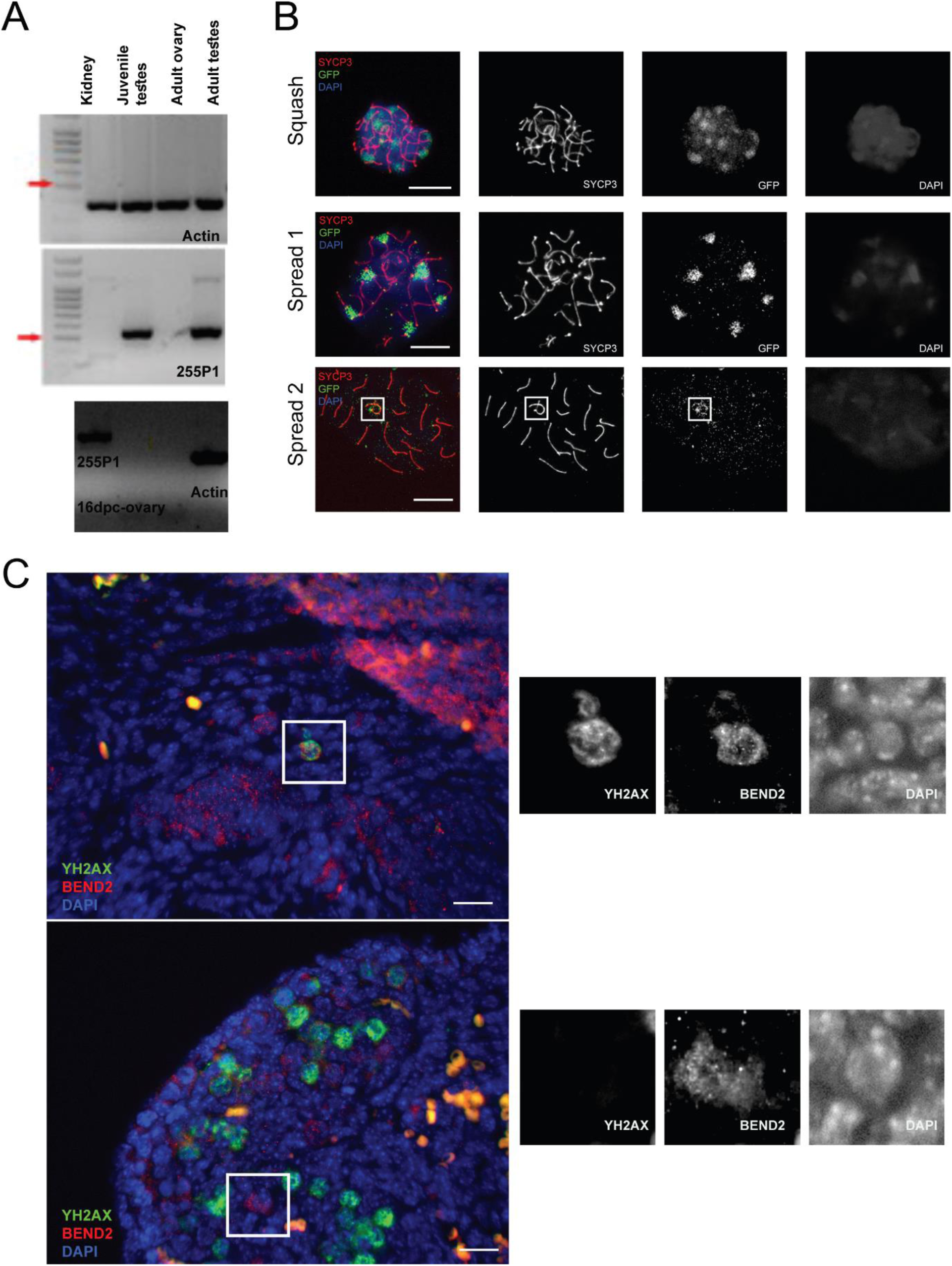
Expression and localization of *Bend2*. (A) Expression of 255P1 in mouse gonads and somatic tissues by RT-PCR. Actin-400bp; 255P1-576bp. (B) Expression pattern of EGFP-255P1 in mouse testis. Representative images of spermatocyte squash and spread stained against SYCP3 and GFP. DNA is counterstained with DAPI (blue). Squash: patch-like signals at the heterochromatic region. Spread 1: cell exhibiting clear foci signals at telomeres along with patch-like signals. Spread 2: cell exhibiting foci signals at the chromosomal telomeres and axes. Signal accumulation at XY chromosomes (white square). Scale bar, 10 µm. (C) BEND2 localization in 16 dpc wild-type mouse ovary. 16 dpc ovary sections were treated with antigen retrieval using Tris-EDTA buffer and stained with BEND2 (red) antibody and ϒH2AX (green) antibody. Strong BEND2 signals were occasionally found in early meiotic prophase oocyte (upper) and oogonia (lower) before DSBs appeared. Scale bar, 20 µm.

**Figure S2.**
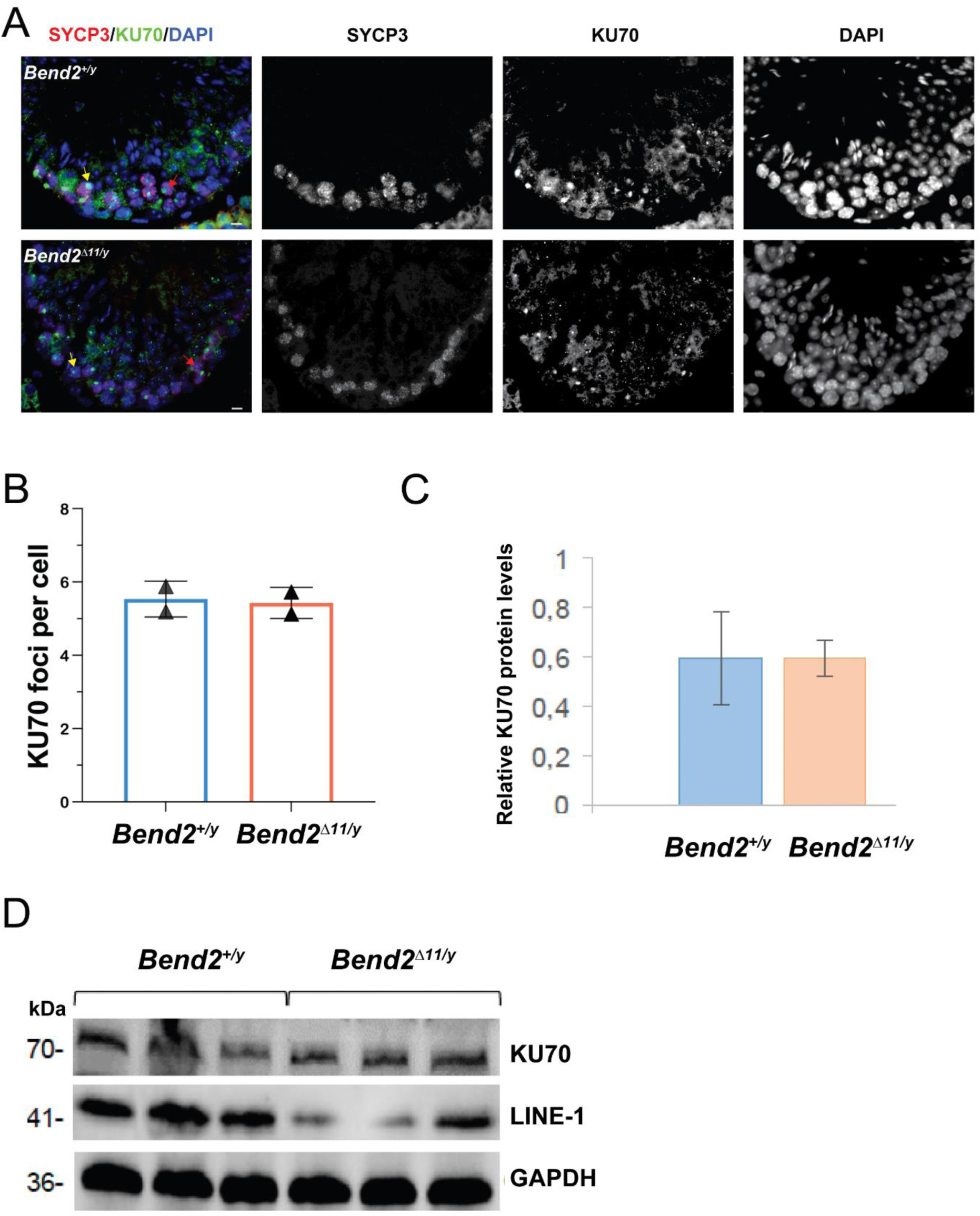
Examination of the NHEJ pathway activity in *Bend2^Δ11/^*^y^ mice. (A) Representative images of Ku70 expression pattern in wild type and *Bend2^Δ11/y^* testis. Scale bar, 40 μm. Yellow arrows indicate the Ku70-positive sex body, and red arrows are two examples of Ku70 foci. (B) Quantification of Ku70 foci in pachytene and diplotene spermatocytes. Foci colocalizing on SYCP3 were counted manually from at least 40 cells per animal. The columns and lines indicate the mean and SD (N=2). (C) Western blot analysis of Ku70 and LINE-1 proteins. (D) Quantification of relative Ku70 protein levels by WB. The columns and lines indicate the mean and SD (N=3).

**Figure S3.**
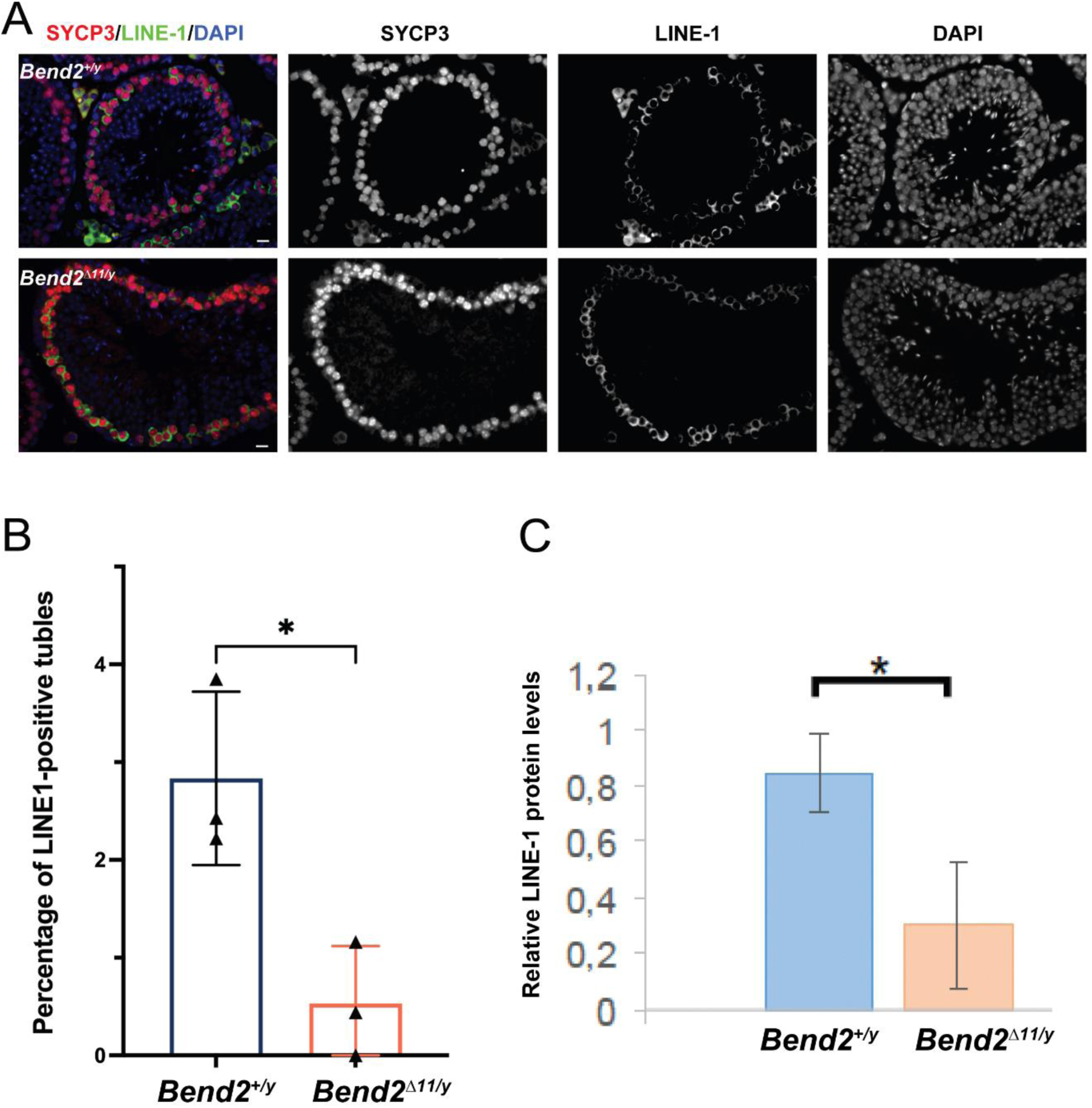
Analysis of LINE-1 retrotransposon expression in *Bend2^Δ11/^*^y^ mice. (A) Representative images of LINE-1 expression pattern in wild type and *Bend2^Δ11/y^* testis. Scale bar, 40 μm. Quantification of LINE-1 positive tubules (B) and relative protein levels in *Bend2^Δ11/y^* animals from Figure S2 (C). The columns and lines indicate the mean and SD (N=3). \**p*=0.046 (B) and *p=0.02 (C) t-test.

**Figure S4.**
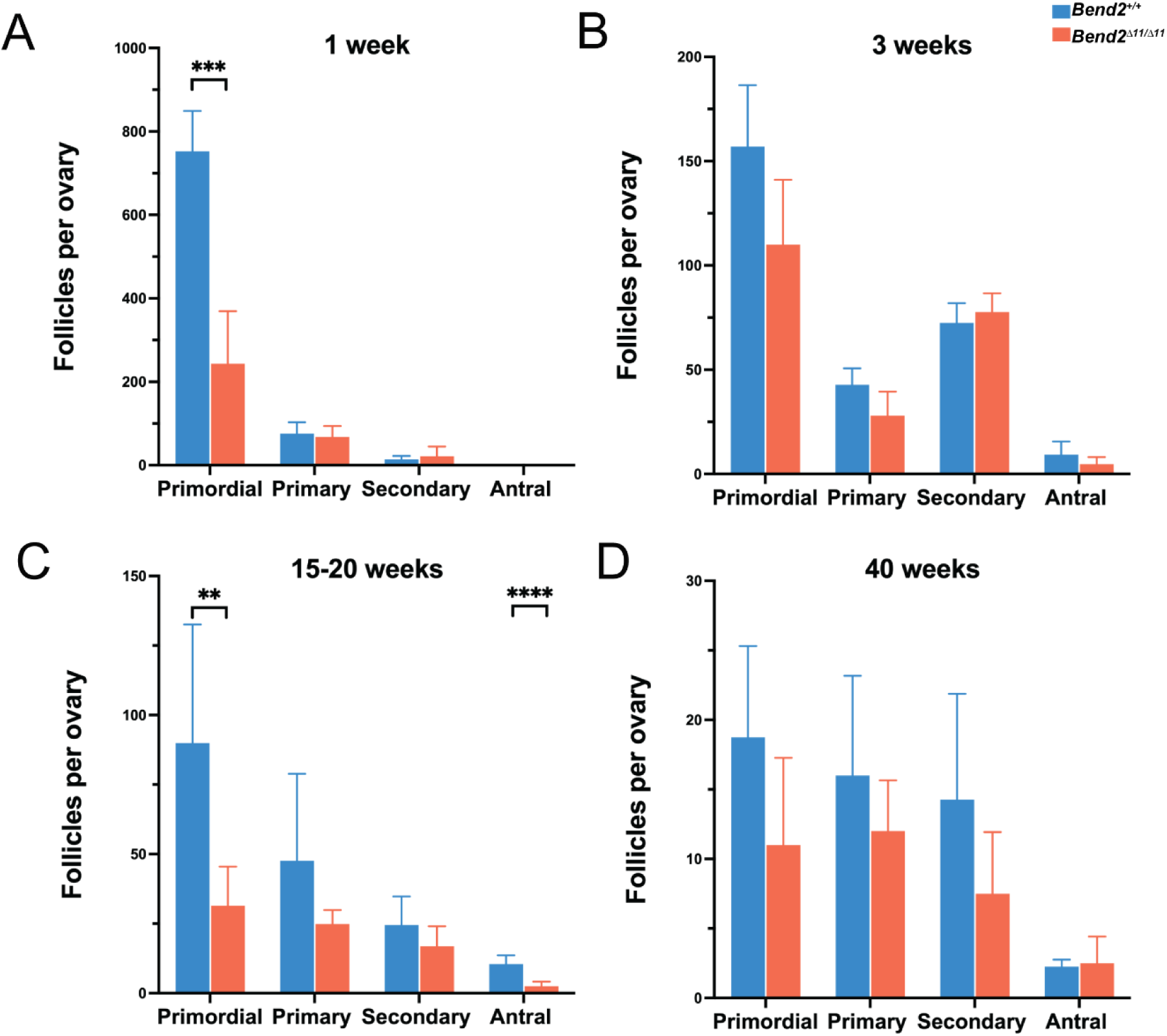
Classification of the type of follicles found in control and *Bend2^Δ11/Δ11^* ovaries. Classification of analyzed follicles from ovaries at the age of 1 week (A), 3 weeks (B), 15-20 weeks (C) and 40 weeks (D). The number of ovaries analyzed per genotype and age is four, except for 15-20 weeks, where eight ovaries were analyzed per genotype. The columns and lines indicate the mean and SD. \*\*\**p*=0.0007 (A); \*\**p*=0.0024, \*\*\*\**p*<0.0001 (C); t-test.

**Figure S5.**
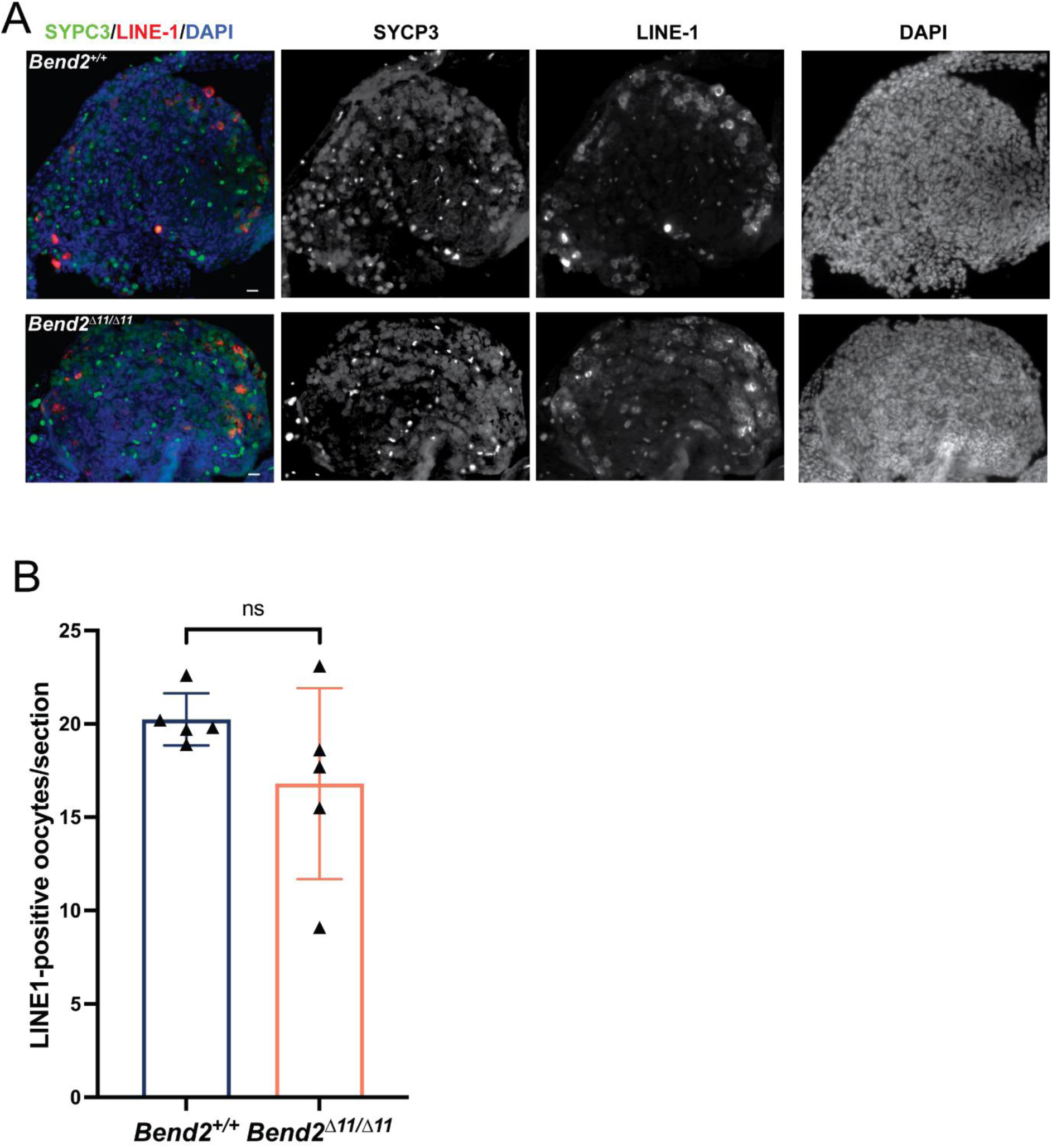
Analysis of LINE-1 retrotransposon expression in *Bend2^Δ11/Δ11^* mice. (A) Representative images of LINE-1 expression pattern in wild type and *Bend2^Δ11/Δ11^* ovaries. Scale bar, 40 μm. (B) Quantification of LINE-1 positive oocytes per section. The columns and lines indicate the mean and SD (N=5), p=0.1851 t-test.

## Supplementary Tables

**Table S1.**
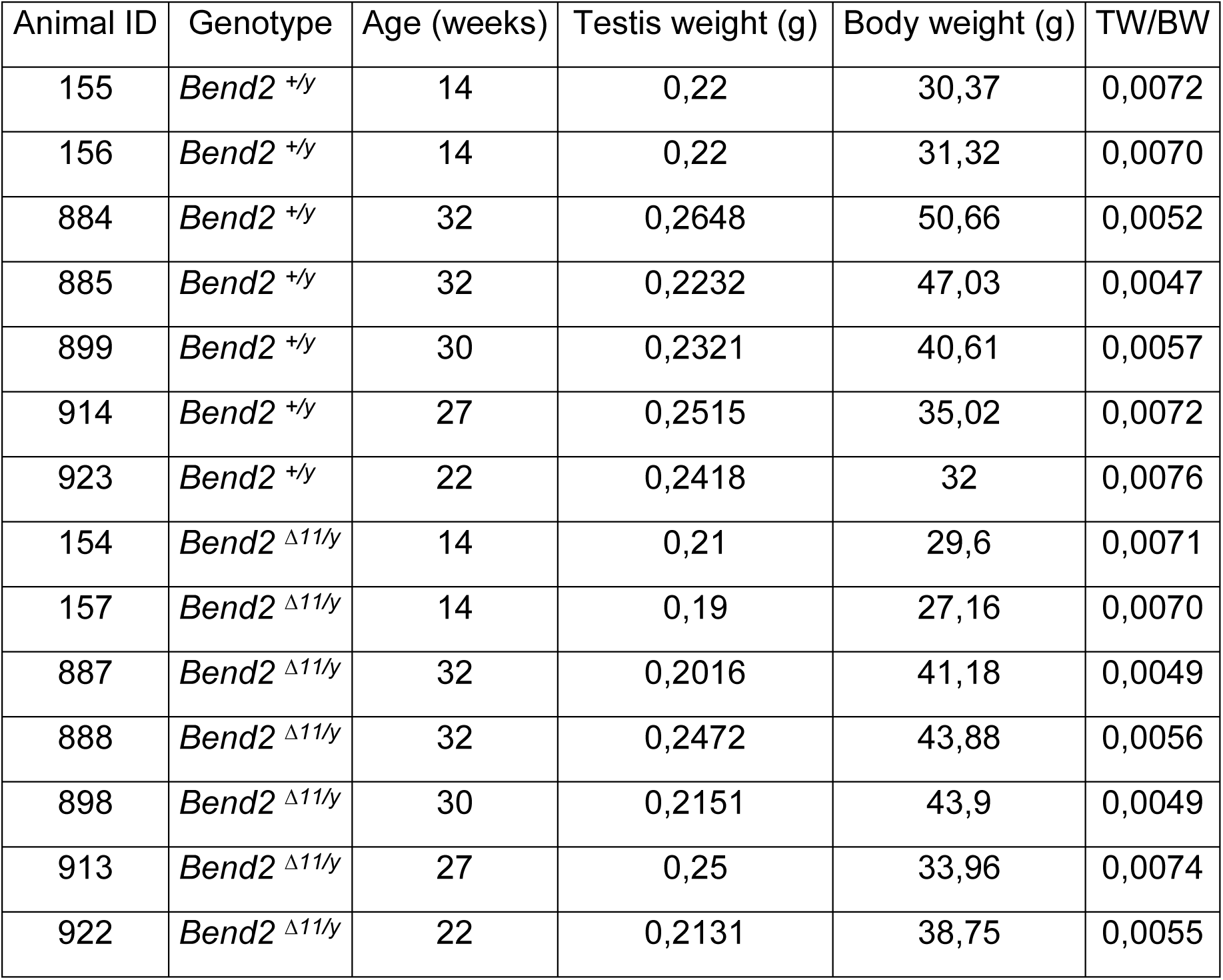
Testis and body weight (TW/BW) ratio of male mice.

## Reference

Abhiman S, Iyer LM, Aravind L. 2008. BEN: A novel domain in chromatin factors and DNA viral proteins. Bioinformatics 24:458. doi:10.1093/BIOINFORMATICS/BTN007

Ahmed EA, Philippens MEP, Kal HB, de Rooij DG, de Boer P. 2010. Genetic probing of homologous recombination and non-homologous end joining during meiotic prophase in irradiated mouse spermatocytes. Mutation Research - Fundamental and Molecular Mechanisms of Mutagenesis 688:12–18. doi:10.1016/j.mrfmmm.2010.02.004

Anderson LK, Reeves A, Webb LM, Ashley T. 1999. Distribution of crossing over on mouse synaptonemal complexes using immunofluorescent localization of MLH1 protein. Genetics 151:1569–1579. doi:10.1093/genetics/151.4.1569

Barchi M, Mahadevaiah S, Di Giacomo M, Baudat F, de Rooij DG, Burgoyne PS, Jasin M, Keeney S. 2005. Surveillance of Different Recombination Defects in Mouse Spermatocytes Yields Distinct Responses despite Elimination at an Identical Developmental Stage. Mol Cell Biol 25:7203–7215. doi:10.1128/mcb.25.16.7203-7215.2005

Baudat F, Manova K, Yuen JP, Jasin M, Keeney S. 2000. Chromosome synapsis defects and sexually dimorphic meiotic progression in mice lacking Spo11. Mol Cell 6:989–998.

Brown MS, Bishop DK. 2015. DNA strand exchange and RecA homologs in meiosis. Cold Spring Harb Perspect Biol 7. doi:10.1101/cshperspect.a016659

Burford A, Mackay A, Popov S, Vinci M, Carvalho D, Clarke M, Izquierdo E, Avery A, Jacques TS, Ingram WJ, Moore AS, Frawley K, Hassall TE, Robertson T, Jones C. 2018. tumor BEN FUSTION The ten-year evolutionary trajectory of a highly recurrent paediatric high grade neuroepithelial tumour with MN1:BEND2 fusion. Sci Rep 8:1–10. doi:10.1038/s41598-018-19389-9

Carofiglio F, Inagaki A, de Vries S, Wassenaar E, Schoenmakers S, Vermeulen C, van Cappellen WA, Sleddens-Linkels E, Grootegoed JA, te Riele HPJ, de Massy B, Baarends WM. 2013. SPO11-Independent DNA Repair Foci and Their Role in Meiotic Silencing. PLoS Genet 9:e1003538. doi:10.1371/journal.pgen.1003538

Dai Q, Ren A, Westholm JO, Serganov AA, Patel DJ, Lai EC. 2013. The BEN domain is a novel sequence-specific DNA-binding domain conserved in neural transcriptional repressors. Genes Dev 27:602. doi:10.1101/GAD.213314.113

Dai Q, Ren A, Westholm JO, Westholm JO, Duan H, Patel DJ, Lai EC. 2015. BEND1 Common and distinct DNA-binding and regulatory activities of the BEN-solo transcription factor family. Genes Dev 29:48–62. doi:10.1101/gad.252122.114

DiGiacomo M, Comazzetto S, Saini H, DeFazio S, Carrieri C, Morgan M, Vasiliauskaite L, Benes V, Enright AJ, O’Carroll D. 2013. Multiple epigenetic mechanisms and the piRNA pathway enforce LINE1 silencing during adult spermatogenesis. Mol Cell 50:601–608. doi:10.1016/J.MOLCEL.2013.04.026

Ding X, Schimenti JC. 2021. Strategies to Identify Genetic Variants Causing Infertility. Trends Mol Med 27:792–806. doi:10.1016/j.molmed.2020.12.008

Enguita-Marruedo A, Martín-Ruiz M, García E, Gil-Fernández A, Parra MT, Viera A, Rufas JS, Page J. 2019. Transition from a meiotic to a somatic-like DNA damage response during the pachytene stage in mouse meiosis. PLoS Genet 15. doi:10.1371/JOURNAL.PGEN.1007439

Garretson A, Dumont BL, Handel MA. 2023. Reproductive genomics of the mouse: implications for human fertility and infertility. Development 150. doi:10.1242/dev.201313

Gerasimova TI, Gdula DA, Gerasimov D V., Simonova O, Corces VG. 1995. A drosophila protein that imparts directionality on a chromatin insulator is an enhancer of position-effect variegation. Cell 82:587–597. doi:10.1016/0092-8674(95)90031-4

Goedecke W, Eijpe M, Offenberg HH, Van Aalderen M, Heyting C. 1999. Mre11 and Ku70 interact in somatic cells, but are differentially expressed in early meiosis. Nat Genet 23:194–198. doi:10.1038/13821

Hinch AG, Becker PW, Li T, Moralli D, Zhang G, Bycroft C, Green C, Keeney S, Shi Q, Davies B, Donnelly P. 2020. The Configuration of RPA, RAD51, and DMC1 Binding in Meiosis Reveals the Nature of Critical Recombination Intermediates. Mol Cell 79:689–701.e10. doi:10.1016/J.MOLCEL.2020.06.015

Huang Y, Roig I. 2023. Genetic control of meiosis surveillance mechanisms in mammals. Front Cell Dev Biol 11. doi:10.3389/FCELL.2023.1127440/FULL

Hunter N. 2017. Oocyte Quality Control: Causes, Mechanisms, and Consequences. Cold Spring Harb Symp Quant Biol 82:235–247. doi:10.1101/sqb.2017.82.035394

Kaul-Ghanekar R, Jalota A, Pavithra L, Tucker P, Chattopadhyay S. 2004. SMAR1 and Cux/CDP modulate chromatin and act as negative regulators of the TCRβ enhancer (Eβ). Nucleic Acids Res 32:4862–4875. doi:10.1093/nar/gkh807

Korutla L, Degnan R, Wang P, Mackler SA. 2007. NAC1, a cocaine-regulated POZ/BTB protein interacts with CoREST. J Neurochem 101:611–618. doi:10.1111/J.1471-4159.2006.04387.X

Korutla L, Wang PJ, Mackler SA. 2005. The POZ/BTB protein NAC1 interacts with two different histone deacetylases in neuronal-like cultures. J Neurochem 94:786–793. doi:10.1111/j.1471-4159.2005.03206.x

Liebe B, Alsheimer M, Höög C, Benavente R, Scherthan H. 2004. Telomere Attachment, Meiotic Chromosome Condensation, Pairing, and Bouquet Stage Duration Are Modified in Spermatocytes Lacking Axial Elements. Mol Biol Cell 15:827–837. doi:10.1091/mbc.E03-07-0524

Ma L, Xie D, Luo M, Lin X, Nie H, Chen J, Gao C, Duo S, Han C. 2022a. Identification and characterization of BEND2 as a key regulator of meiosis during mouse spermatogenesis. Sci Adv 8. doi:10.1126/SCIADV.ABN1606

Ma L, Xie D, Luo M, Lin X, Nie H, Chen J, Gao C, Duo S, Han C. 2022b. Identification and characterization of BEND2 as a key regulator of meiosis during mouse spermatogenesis. Sci Adv 8. doi:10.1126/sciadv.abn1606

Malcher A, Stokowy T, Berman A, Olszewska M, Jedrzejczak P, Sielski D, Nowakowski A, Rozwadowska N, Yatsenko AN, Kurpisz MK. 2022. Whole-genome sequencing identifies new candidate genes for nonobstructive azoospermia. Andrology 10:1605–1624. doi:10.1111/ANDR.13269

Malki S, van der Heijden GW, O’Donnell KA, Martin SL, Bortvin A. 2019. A Role for Retrotransposon LINE-1 in Fetal Oocyte Attrition in Mice. Dev Cell 51:658. doi:10.1016/j.devcel.2019.11.011

Malki S, vanderHeijden GW, O’Donnell KA, Martin SL, Bortvin A. 2014. A Role for retrotransposon LINE-1 in fetal oocyte attrition in mice. Dev Cell 29:521–533. doi:10.1016/j.devcel.2014.04.027

Marcet Ortega Marina, Roig I (Ignasi), Universitat Autònoma de Barcelona. Departament de Biologia Cel·lular de F i d’Immunologia. 2016. Surveillance mechanisms in mammalian meiosis. TDX (Tesis Doctorals en Xarxa).

Martínez-Marchal A, Huang Y, Guillot-Ferriols MT, Ferrer-Roda M, Guixé A, Garcia-Caldés M, Roig I. 2020. The DNA damage response is required for oocyte cyst breakdown and follicle formation in mice. PLoS Genet 16:e1009067. doi:10.1371/journal.pgen.1009067

Moens PB, Kolas NK, Tarsounas M, Marcon E, Cohen PE, Spyropoulos B. 2002. The time course and chromosomal localization of recombination-related proteins at meiosis in the mouse are compatible with models that can resolve the early DNA-DNA interactions without reciprocal recombination. J Cell Sci 115:1611–1622. doi:10.1242/JCS.115.8.1611

Nègre N, Brown CD, Shah PK, Kheradpour P, Morrison CA, Henikoff JG, Feng X, Ahmad K, Russell S, White RAH, Stein L, Henikoff S, Kellis M, White KP. 2010. A Comprehensive Map of Insulator Elements for the Drosophila Genome. PLoS Genet 6:e1000814. doi:10.1371/JOURNAL.PGEN.1000814

Pacheco S, Marcet-Ortega M, Lange J, Jasin M, Keeney S, Roig I. 2015. The ATM Signaling Cascade Promotes Recombination-Dependent Pachytene Arrest in Mouse Spermatocytes. PLoS Genet 11:1–27. doi:10.1371/journal.pgen.1005017

Pittman DL, Cobb J, Schimenti KJ, Wilson LA, Cooper DM, Brignull E, Handel MA, Schimenti JC. 1998. Meiotic prophase arrest with failure of chromosome synapsis in mice deficient for Dmc1, a germline-specific RecA homolog. Mol Cell 1:697–705.

Rampalli S, Pavithra L, Bhatt A, Kundu TK, Chattopadhyay S. 2005. Tumor Suppressor SMAR1 Mediates Cyclin D1 Repression by Recruitment of the SIN3/Histone Deacetylase 1 Complex. Mol Cell Biol 25:8415–8429. doi:10.1128/mcb.25.19.8415-8429.2005

Riera-Escamilla A, Enguita-Marruedo A, Moreno-Mendoza D, Chianese C, Sleddens-Linkels E, Contini E, Benelli M, Natali A, Colpi GM, Ruiz-Castañé E, Maggi M, Baarends WM, Krausz C. 2019. Sequencing of a “mouse azoospermia” gene panel in azoospermic men: Identification of RNF212 and STAG3 mutations as novel genetic causes of meiotic arrest. Human Reproduction 34:978–988. doi:10.1093/humrep/dez042

Roig I, Liebe B, Egozcue J, Cabero L, Garcia M, Scherthan H. 2004. Female-specific features of recombinational double-stranded DNA repair in relation to synapsis and telomere dynamics in human oocytes. Chromosoma 113:22–33. doi:10.1007/s00412-004-0290-8

Rossetti R, Ferrari I, Bonomi M, Persani L. 2017. Genetics of primary ovarian insufficiency. Clin Genet. doi:10.1111/cge.12921

Ruth KS, Day FR, Hussain J, Martínez-Marchal A, Aiken CE, Azad A, Thompson DJ, Knoblochova L, Abe H, Tarry-Adkins JL, Gonzalez JM, Fontanillas P, Claringbould A, Bakker OB, Sulem P, Walters RG, Terao C, Turon S, Horikoshi M, Lin K, Onland-Moret NC, Sankar A, Hertz EPT, Timshel PN, Shukla V, Borup R, Olsen KW, Aguilera P, Ferrer-Roda M, Huang Y, Stankovic S, Timmers PRHJ, Ahearn TU, Alizadeh BZ, Naderi E, Andrulis IL, Arnold AM, Aronson KJ, Augustinsson A, Bandinelli S, Barbieri CM, Beaumont RN, Becher H, Beckmann MW, Benonisdottir S, Bergmann S, Bochud M, Boerwinkle E, Bojesen SE, Bolla MK, Boomsma DI, Bowker N, Brody JA, Broer L, Buring JE, Campbell A, Campbell H, Castelao JE, Catamo E, Chanock SJ, Chenevix-Trench G, Ciullo M, Corre T, Couch FJ, Cox A, Crisponi L, Cross SS, Cucca F, Czene K, Smith GD, de Geus EJCN, de Mutsert R, De Vivo I, Demerath EW, Dennis J, Dunning AM, Dwek M, Eriksson M, Esko T, Fasching PA, Faul JD, Ferrucci L, Franceschini N, Frayling TM, Gago-Dominguez M, Mezzavilla M, García-Closas M, Gieger C, Giles GG, Grallert H, Gudbjartsson DF, Gudnason V, Guénel P, Haiman CA, Håkansson N, Hall P, Hayward C, He C, He W, Heiss G, Høffding MK, Hopper JL, Hottenga JJ, Hu F, Hunter D, Ikram MA, Jackson RD, Joaquim MDR, John EM, Joshi PK, Karasik D, Kardia SLR, Kartsonaki C, Karlsson R, Kitahara CM, Kolcic I, Kooperberg C, Kraft P, Kurian AW, Kutalik Z, La Bianca M, LaChance G, Langenberg C, Launer LJ, Laven JSE, Lawlor DA, Le Marchand L, Li J, Lindblom A, Lindstrom S, Lindstrom T, Linet M, Liu YM, Liu S, Luan J, Mägi R, Magnusson PKE, Mangino M, Mannermaa A, Marco B, Marten J, Martin NG, Mbarek H, McKnight B, Medland SE, Meisinger C, Meitinger T, Menni C, Metspalu A, Milani L, Milne RL, Montgomery GW, Mook-Kanamori DO, Mulas A, Mulligan AM, Murray Alison, Nalls MA, Newman A, Noordam R, Nutile T, Nyholt DR, Olshan AF, Olsson H, Painter JN, Patel A V., Pedersen NL, Perjakova N, Peters A, Peters U, Pharoah PDP, Polasek O, Porcu E, Psaty BM, Rahman I, Rennert G, Rennert HS, Ridker PM, Ring SM, Robino A, Rose LM, Rosendaal FR, Rossouw J, Rudan I, Rueedi R, Ruggiero D, Sala CF, Saloustros E, Sandler DP, Sanna S, Sawyer EJ, Sarnowski C, Schlessinger D, Schmidt MK, Schoemaker MJ, Schraut KE, Scott C, Shekari S, Shrikhande A, Smith A V., Smith BH, Smith JA, Sorice R, Southey MC, Spector TD, Spinelli JJ, Stampfer M, Stöckl D, van Meurs JBJ, Strauch K, Styrkarsdottir U, Swerdlow AJ, Tanaka T, Teras LR, Teumer A, Þorsteinsdottir U, Timpson NJ, Toniolo D, Traglia M, Troester MA, Truong T, Tyrrell J, Uitterlinden AG, Ulivi S, Vachon CM, Vitart V, Völker U, Vollenweider P, Völzke H, Wang Q, Wareham NJ, Weinberg CR, Weir DR, Wilcox AN, van Dijk KW, Willemsen G, Wilson JF, Wolffenbuttel BHR, Wolk A, Wood AR, Zhao W, Zygmunt M, Chen Z, Li L, Franke L, Burgess S, Deelen P, Pers TH, Grøndahl ML, Andersen CY, Pujol A, Lopez-Contreras AJ, Daniel JA, Stefansson K, Chang-Claude J, van der Schouw YT, Lunetta KL, Chasman DI, Easton DF, Visser JA, Ozanne SE, Namekawa SH, Solc P, Murabito JM, Ong KK, Hoffmann ER, Murray Anna, Roig I, Perry JRB. 2021. Genetic insights into biological mechanisms governing human ovarian ageing. Nature 596:393–397. doi:10.1038/S41586-021-03779-7

Sathyan KM, Shen Z, Tripathi V, Prasanth K V., Prasanth SG. 2011. A BEN-domain-containing protein associates with heterochromatin and represses transcription. J Cell Sci 124:3149–3163. doi:10.1242/jcs.086603

Scarpa A, Chang DK, Nones K, Corbo V, Patch AM, Bailey P, Lawlor RT, Johns AL, Miller DK, Mafficini A, Rusev B, Scardoni M, Antonello D, Barbi S, Sikora KO, Cingarlini S, Vicentini C, McKay S, Quinn MCJ, Bruxner TJC, Christ AN, Harliwong I, Idrisoglu S, McLean S, Nourse C, Nourbakhsh E, Wilson PJ, Anderson MJ, Fink JL, Newell F, Waddell Nick, Holmes O, Kazakoff SH, Leonard C, Wood S, Xu Q, Nagaraj SH, Amato E, Dalai I, Bersani S, Cataldo I, Dei Tos AP, Capelli P, Davì MV, Landoni L, Malpaga A, Miotto M, Whitehall VLJ, Leggett BA, Harris JL, Harris J, Jones MD, Humphris J, Chantrill LA, Chin V, Nagrial AM, Pajic M, Scarlett CJ, Pinho A, Rooman I, Toon C, Wu J, Pinese M, Cowley M, Barbour A, Mawson A, Humphrey ES, Colvin EK, Chou A, Lovell JA, Jamieson NB, Duthie F, Gingras MC, Fisher WE, Dagg RA, Lau LMS, Lee M, Pickett HA, Reddel RR, Samra JS, Kench JG, Merrett ND, Epari K, Nguyen NQ, Zeps N, Falconi M, Simbolo M, Butturini G, Van Buren G, Partelli S, Fassan M, Khanna KK, Gill AJ, Wheeler DA, Gibbs RA, Musgrove EA, Bassi C, Tortora G, Pederzoli P, Pearson J V., Waddell Nicola, Biankin A V., Grimmond SM. 2017. Whole-genome landscape of pancreatic neuroendocrine tumours. Nature 543:65–71. doi:10.1038/nature21063

Shibuya H, Morimoto A, Watanabe Y. 2014. The Dissection of Meiotic Chromosome Movement in Mice Using an In Vivo Electroporation Technique. PLoS Genet 10:e1004821. doi:10.1371/journal.pgen.1004821

Soper SFC, van der Heijden GW, Hardiman TC, Goodheart M, Martin SL, de Boer P, Bortvin A. 2008. Mouse Maelstrom, a component of nuage, is essential for spermatogenesis and transposon repression in meiosis. Dev Cell 15:285. doi:10.1016/J.DEVCEL.2008.05.015

Sturm D, Orr BA, Toprak UH, Hovestadt V, Jones DTW, Capper D, Sill M, Buchhalter I, Northcott PA, Leis I, Ryzhova M, Koelsche C, Pfaff E, Allen SJ, Balasubramanian G, Worst BC, Pajtler KW, Brabetz S, Johann PD, Sahm F, Reimand J, Mackay A, Carvalho DM, Remke M, Phillips JJ, Perry A, Cowdrey C, Drissi R, Fouladi M, Giangaspero F, Łastowska M, Grajkowska W, Scheurlen W, Pietsch T, Hagel C, Gojo J, Lötsch D, Berger W, Slavc I, Haberler C, Jouvet A, Holm S, Hofer S, Prinz M, Keohane C, Fried I, Mawrin C, Scheie D, Mobley BC, Schniederjan MJ, Santi M, Buccoliero AM, Dahiya S, Kramm CM, Von Bueren AO, Von Hoff K, Rutkowski S, Herold-Mende C, Frühwald MC, Milde T, Hasselblatt M, Wesseling P, Rößler J, Schüller U, Ebinger M, Schittenhelm J, Frank S, Grobholz R, Vajtai I, Hans V, Schneppenheim R, Zitterbart K, Collins VP, Aronica E, Varlet P, Puget S, Dufour C, Grill J, Figarella-Branger D, Wolter M, Schuhmann MU, Shalaby T, Grotzer M, Van Meter T, Monoranu CM, Felsberg J, Reifenberger G, Snuderl M, Forrester LA, Koster J, Versteeg R, Volckmann R, Van Sluis P, Wolf S, Mikkelsen T, Gajjar A, Aldape K, Moore AS, Taylor MD, Jones C, Jabado N, Karajannis MA, Eils R, Schlesner M, Lichter P, Von Deimling A, Pfister SM, Ellison DW, Korshunov A, Kool M. 2016. New Brain Tumor Entities Emerge from Molecular Classification of CNS-PNETs. Cell 164:1060–1072. doi:10.1016/J.CELL.2016.01.015

Tharp ME, Malki S, Bortvin A. 2020. Maximizing the ovarian reserve in mice by evading LINE-1 genotoxicity. Nature Communications 2020 11:1 11:1–13. doi:10.1038/s41467-019-14055-8

Wang PJ. 2017. Tracking LINE1 retrotransposition in the germline. Proc Natl Acad Sci U S A 114:7194–7196. doi:10.1073/PNAS.1709067114

Williamson LM, Steel M, Grewal JK, Thibodeau ML, Zhao EY, Loree JM, Yang KC, Gorski SM, Mungall AJ, Mungall KL, Moore RA, Marra MA, Laskin J, Renouf DJ, Schaeffer DF, Jones SJM. 2019. tumor BEN FUSTION Genomic characterization of a welldifferentiated grade 3 pancreatic neuroendocrine tumor. Cold Spring Harb Mol Case Stud 5:1–14. doi:10.1101/mcs.a003814

Xuan C, Wang Q, Han X, Duan Y, Li L, Shi L, Wang Y, Shan L, Yao Z, Shang Y. 2012. RBB, a novel transcription repressor, represses the transcription of HDM2 oncogene. Oncogene 2013 32:32 32:3711–3721. doi:10.1038/onc.2012.386

Zickler D, Kleckner N. 1999. Meiotic chromosomes: Integrating structure and function. Annu Rev Genet. doi:10.1146/annurev.genet.33.1.603

